# Lab on a Capillary: Instrument-Free Compartmentalization Using Photopatterned Hydrogel Rings Embedded inside a Glass Capillary for Amplified Bioassays

**DOI:** 10.64898/2026.07.08.737077

**Authors:** Yimin Yang, Muhammad Usman Akhtar, Mehmet Akif Sahin, Yibo Huang, Liusu Wang, Xinpei Song, Ghulam Destgeer

## Abstract

Sensitive and low-cost protein biomarker detection is critical for disease diagnosis. Advanced microfluidic systems can generate miniature reaction compartments for a high-sensitivity assay. However, these platforms often require external instruments, skilled operators, and complex setups. Here, we develop a Lab on a Capillary (LabCap) platform that integrates photopatterned hydrogel rings within a glass capillary using a reconfigurable stop-flow lithography system. During sample loading and unloading steps, nanoliter-scale aqueous droplets (torodrops) are spontaneously formed around the hydrogel rings, creating isolated reaction compartments without the need for external instruments or an immiscible oil phase. The LabCap platform enables quantitative detection of clinically relevant biomarkers, including C-reactive protein (CRP) and N-terminal pro-B-type natriuretic peptide (NT-proBNP). By adjusting the incubation protocol, assay speed and sensitivity can be tuned to meet different analytical requirements. A periodic medium exchange protocol enables biomarker detection at concentrations as low as 1 ng/mL, whereas prolonged static incubation extends detection to 0.1 ng/mL. In addition, LabCap offers practical advantages, including low fabrication cost (<€1 per device), low reagent consumption (<100 μL per assay step), and minimal wash-buffer usage (1 mL). These results demonstrate that LabCap is a simple, cost-effective, and versatile platform for biomarker detection.

## 1. Introduction

Affordable and sensitive protein biomarker detection is essential for disease diagnosis and therapeutic monitoring, particularly in resource-limited regions.^1,2^ Achieving high analytical sensitivity often requires confining target analytes within small reaction volumes to increase effective concentration and accelerate reaction kinetics. Microfluidic droplets and microwell arrays have been widely employed for biomolecule detection^3–5^ and single-cell analysis.^6–8^ Despite their excellent analytical performance, these platforms generally depend on cleanroom fabrication, external instrumentation, and skilled operation, limiting their accessibility outside specialized laboratories. Consequently, there remains a critical need for compartmentalized analytical platforms that combine high sensitivity with simple and low-cost operation.

Hydrogel-based biosensing strategies have attracted considerable attention due to their high water content, tunable chemistry, and porous polymer networks that facilitate efficient biomolecular diffusion and high-capacity binding sites for target capture.^9^ To improve detection sensitivity and minimize signal crosstalk, microfluidic devices have been developed to isolate hydrogel particles within individual microcompartments.^10–13^ However, the sophisticated device architectures and operational complexity required for particle trapping and compartmentalization limit their practical application. To simplify the compartmentalization process, particle-templated droplet strategies have been proposed,^14^ where hydrogel particles act as physical templates to form isolated aqueous microcompartments upon mixing with oil via simple laboratory operations such as pipetting or vortexing.^15–18^ In these systems, aqueous solutions were confined either within cavities of the hydrogel particles^19,20^ or as thin shells surrounding the particles,^14,17^ locally concentrating reaction product and enhancing detectable signals, while the immiscible oil suppresses crosstalk between neighboring compartments. Despite eliminating the need for microfluidic equipment during assay operation, hydrogel particle fabrication typically still requires specialized microfluidic devices and lab apparatus. Additionally, the use of surfactants to stabilize droplets raises concerns regarding signal leakage.^21,22^ Paper-based analytical devices offer an alternative route toward instrument-free operation by integrating hydrogels within cellulose substrates and utilizing capillary wicking for fluid transport.^23,24^ While these paper-hydrogel hybrid platforms are inexpensive and easy to operate, the heterogeneous pore structure of paper makes it difficult to precisely control hydrogel geometry and reaction volume, limiting reproducibility and quantitative performance.

To overcome these limitations, capillary-based platforms offer a promising alternative for compartmentalized bioassay. Glass capillaries are attractive because they combine autonomous capillary-driven fluid transport with optical transparency, low cost, ease of handling and long-term storage.^25,26^ Furthermore, their well-defined geometry enables precise control over reaction volumes and hydrogel structures. Previous studies have demonstrated capillary-based enzyme-linked immunosorbent assay (ELISA), in which antibodies are directly immobilized on the inner surface of a glass capillary through chemical coupling reaction such as EDC/NHS chemistry.^27,28^ These systems simplify reagent handling and eliminate the need for external pumping equipment. However, uniform surface functionalization transforms the entire capillary into a single continuous reaction zone, inherently limiting spatial compartmentalization and localized signal amplification. Introducing spatially defined hydrogel structures within a capillary provides an attractive strategy for creating multiple discrete reaction compartments while preserving the simplicity of capillary-driven operation.

Herein, we present an oil-free Lab on a Capillary (LabCap) platform that requires no external instruments during assay operation. The LabCap platform integrates photopatterned hydrogel rings inside a glass capillary using a reconfigurable stop-flow lithography system. The platform consists of annular hydrogel arrays that serve as three-dimensional reaction scaffolds and can be functionalized via streptavidin-biotin chemistry to immobilize capture antibody (**Figure 1**). The anchored hydrogel rings form oil- and surfactant-free, spatially addressable torodrops, whereas the pre-functionalized glass capillaries can be stored for later use, thereby significantly reducing the assay time. During assay operation, reagents are introduced sequentially by capillary action to perform sandwich immunoassays within hydrogel ring. Following immunocomplex formation, removal of the bulk solution induces spontaneous liquid breakup, generating nanoliter-scale toroidal droplets (torodrops) around each hydrogel ring. These confined reaction volumes localize enzymatic signal generation and enhance fluorescence readout. Using the LabCap platform, we demonstrate sensitive and quantitative detection of clinically relevant protein biomarkers, including C-reactive protein (CRP) and N-terminal pro-B-type natriuretic peptide (NT-proBNP). The LabCap platform provides a simple, low-cost, and versatile approach for biomarker detection.

**Figure 1.**
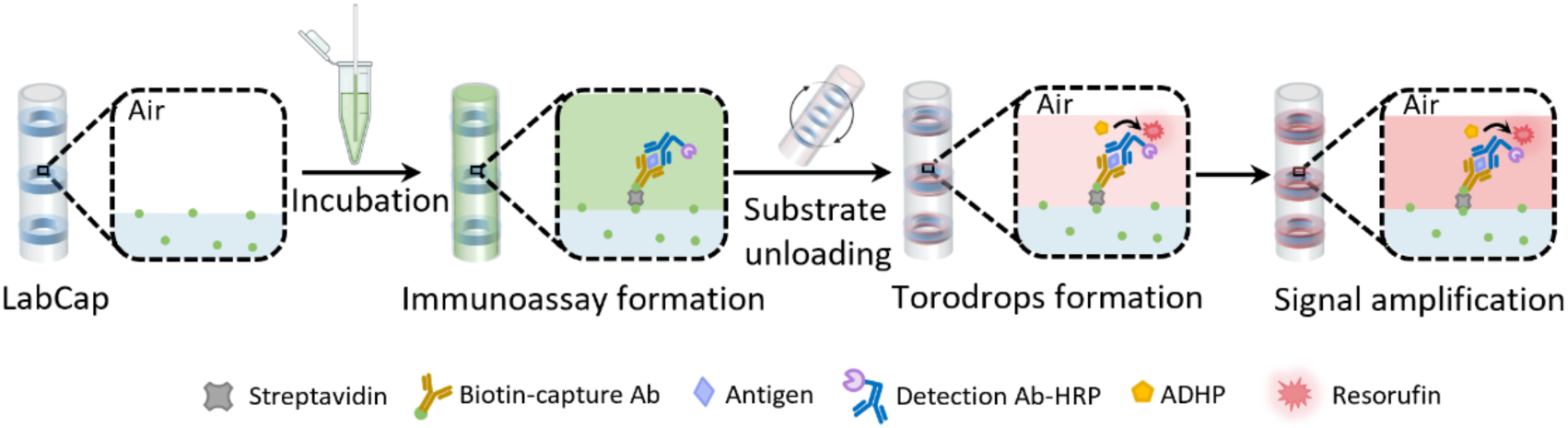
Schematic illustration of the Lab on a Capillary (LabCap) platform for compartmentalized amplified bioassays. Biotinylated hydrogel rings photopatterned inside a glass capillary are functionalized with capture antibodies and used for target binding. Following sandwich immunoassay formation, removal of the bulk solution generates nanoliter-scale torodrop around each hydrogel ring. Enzymatic conversion of ADHP substrate to resorufin within the confined compartments produces amplified fluorescence signals for biomarker detection.

## 2. Results and discussion

### 2.1 Stop-flow lithography for embedding hydrogel rings inside a glass capillary

To generate spatially discrete compartments inside a glass capillary, hydrogel rings were patterned and immobilized along the inner wall of the capillary using stop-flow lithography. Stable anchoring of the hydrogel structures was achieved by first treating the capillary with NaOH and subsequently silanizing the inner surface with 3-(trimethoxysilyl)propyl methacrylate (TMSPMA), introducing polymerizable methacrylate groups for covalent attachment of PEGDA hydrogel (**Figure 2a**). Hydrogel ring fabrication was performed using a reconfigurable stop-flow lithography setup consisting of a 3D-printed concentric microfluidic device coupled to a detachable glass capillary. A photocurable outer stream containing poly(ethylene glycol) diacrylate (PEGDA) and photoinitiator (PI), and an inert inner stream containing poly(ethylene glycol) (PEG) and PI, were introduced through concentric channels to generate a coaxial flow (Figure 2b). After the flow was established within the capillary, it was temporarily halted and exposed to patterned UV illumination through a photomask (Figure 2c). The PEGDA phase was selectively crosslinked to form annular hydrogel structure covalently anchored to the capillary wall, while the PEG core remained uncrosslinked and was subsequently flushed out. The detachable capillary configuration enabled repeated fabrication using a single 3D-printed device, resulting in a material cost of < €1 per device (**Table S1**).

**Figure 2.**
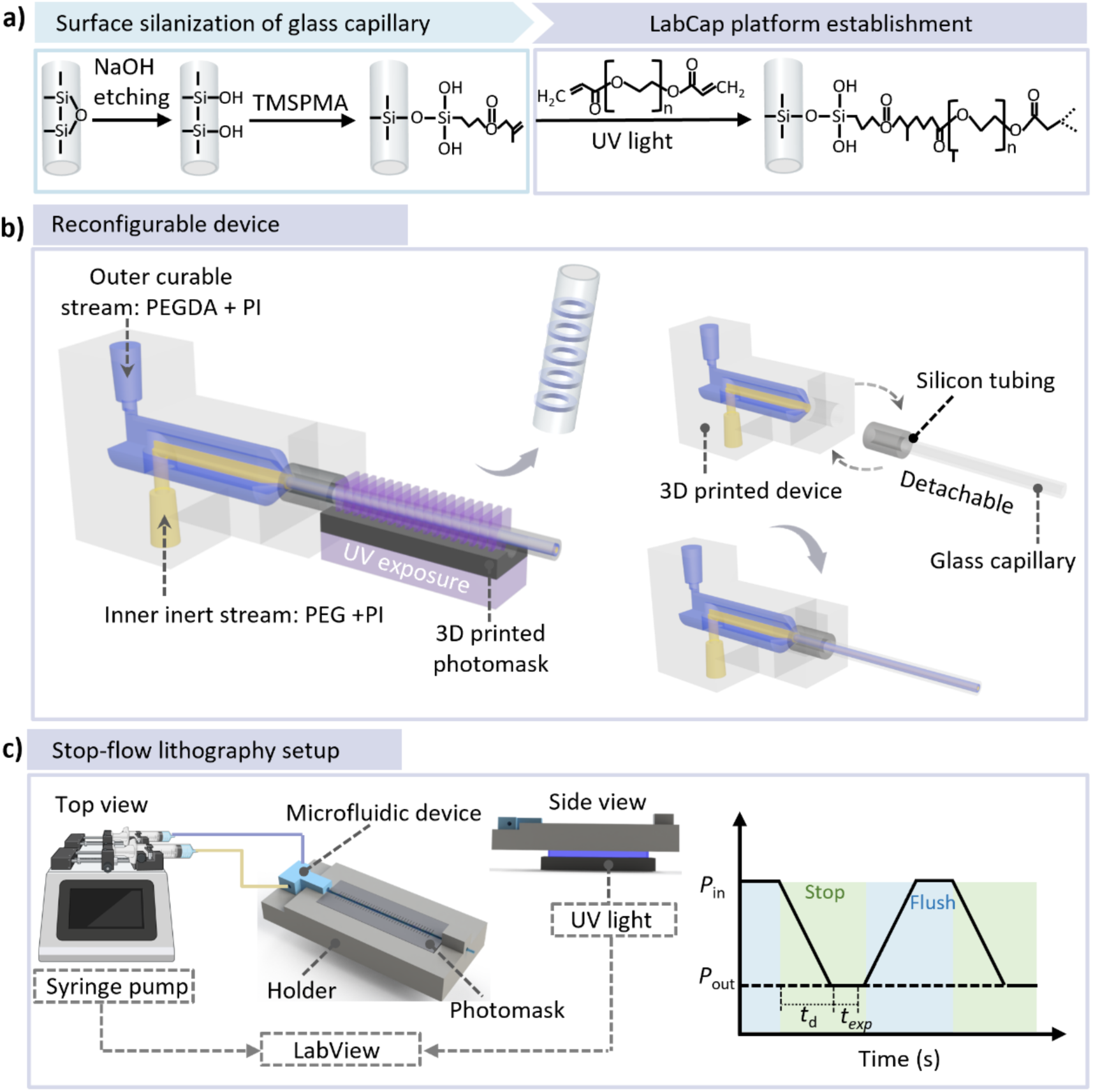
Stop-flow lithography for embedding hydrogel rings inside a glass capillary. (a) Surface modification of the glass capillary. The inner wall of the capillary was activated by NaOH etching and silanized with 3-(trimethoxysilyl)propyl methacrylate (TMSPMA) to introduce polymerizable methacrylate groups for covalent anchoring of PEGDA hydrogels. (b) Schematic of the reconfigurable 3D printed concentric microfluidic device coupled to a detachable glass capillary via a compliant silicone tubing interface. Independent inner and outer inlets deliver inert PEG and photocurable PEGDA streams, respectively, to generate coaxial flow. (c) Stop-flow lithography process. The coaxial flow was pumped into a glass capillary, generating an inlet pressure (*P*_in_), after a short delay time (*t_d_*), the flow was stopped, allowing the pressure inside the capillary to equilibrate to atmospheric pressure (*P*_out_). The stationary flow was then exposed to patterned UV through a photomask for a defined exposure time (*t*_exp_) selectively crosslinking hydrogel rings covalently immobilized on the capillary wall. Schematics were partially created with BioRender.

### 2.2. Instrument-free formation of torodrops using capillary action

The glass capillary embedded with hydrogel rings was employed to generate miniature aqueous compartments following sequential washing with ethanol and water (**Figure 3a**). To visualize compartment formation, the hydrogel rings were labeled with red-emitting fluorescent dye, while green fluorescein solution was used for torodrops formation. Upon sample loading by capillary action, the porous hydrogel rings became fully hydrated and saturated with the fluorescein solution. To initiate compartment formation, the capillary was inverted and briefly contacted with an absorbent substrate (e.g., tissue paper) to remove the excess liquid. As the liquid receded, the air-liquid meniscus propagated along the capillary. When the receding contact line encountered a hydrogel ring, it became pinned at the hydrogel boundary due to the combined effects of the topographical discontinuity and highly hydrophilic hydrogel surface. Continued liquid withdrawal stretched the pinned liquid into a thin liquid bridge, which subsequently underwent capillary instability and ruptured. The residual liquid surrounding the hydrogel ring then relaxed into a stable toroidal microdroplet while remaining anchored to the hydrogel structure. These confined aqueous volumes, referred to as torodrops, serve as nanoliter-scale microreactors for subsequent biochemical reactions. Importantly, torodrops formation occurs spontaneously without the need for external pumping systems, active flow control, or an immiscible oil phase, enabling simple and instrument-free compartmentalization within the capillary.

**Figure 3.**
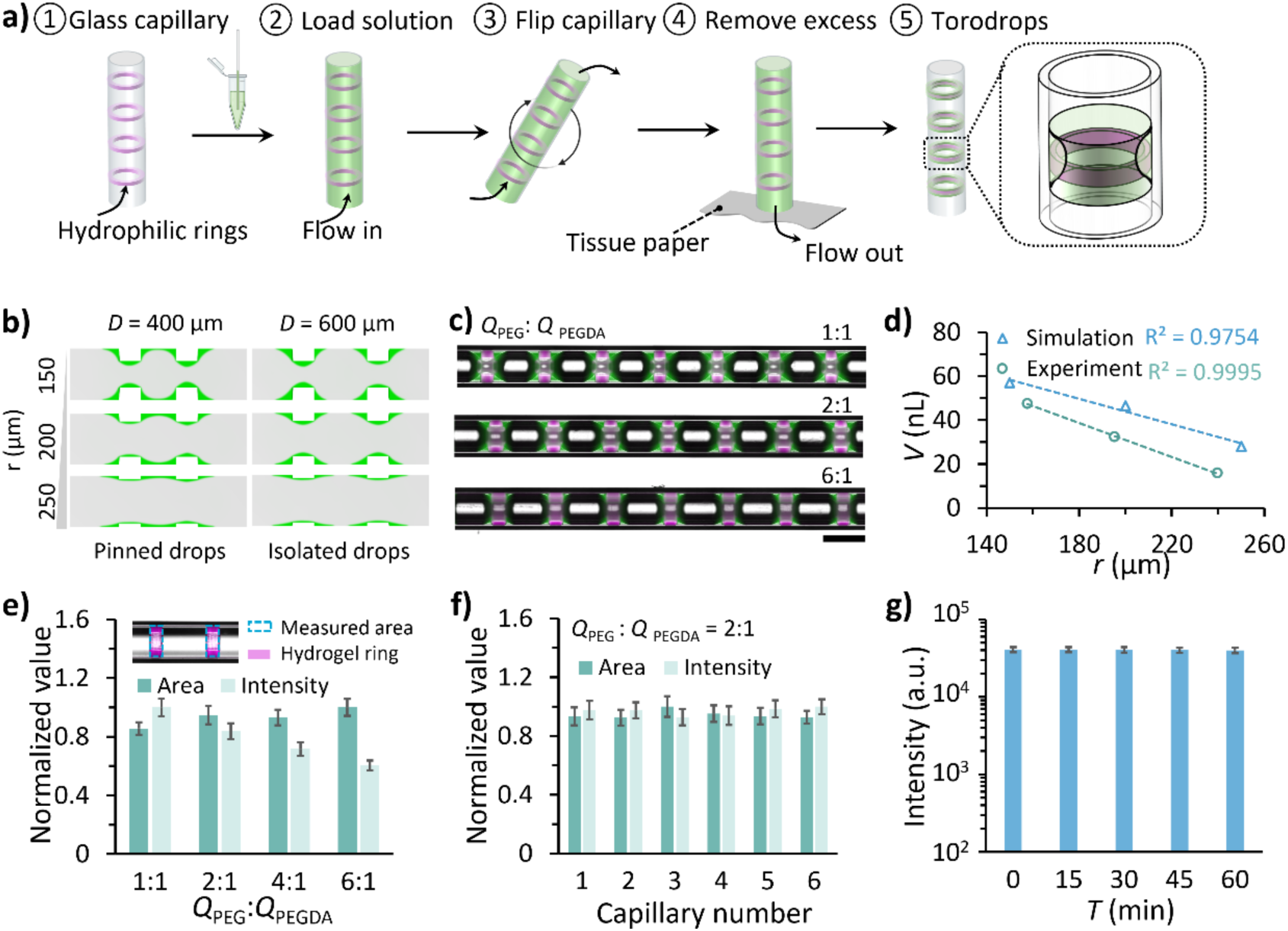
Characterization of LabCap platform and torodrop formation. (a) Schematic illustration of torodrops formation following aqueous solution loading and unloading. (b) Numerical models of the torodrops formation within capillary as a function of the inner radius of hydrogel rings (*r*) and distance between adjacent hydrogel rings (*D*). (c) Merged images of bright-field and fluorescence channels showing torodrops formed around fluorescently labeled hydrogel rings fabricated at different flow rate ratios of PEG and PEGDA. Hydrogel rings were patterned at 1 mm intervals. The scale bar represents 1 mm. (d) Experimental estimated and numerically predicted torodrop volumes as a function of *r*. (e) The normalized area and fluorescence intensity of hydrogel rings fabricated with four different *Q*_PEG_:*Q*_PEGDA_ from 1:1 to 6:1. The fluorescence intensity and area of hydrogel rings were normalized against the overall maximum intensity and area. The inset illustrates the measured region used for fluorescence intensity and area quantification. (f) The normalized area and fluorescence intensity of hydrogel rings fabricated in six batches at *Q*_PEG_:*Q*_PEGDA_ of 2:1. (g) The fluorescence intensity of hydrogel rings excluding the two rings closest to the capillary ends after 1 h incubation. Error bars represent standard deviation.

### 2.3 Characterization of LabCap

To understand the equilibrium isolation and volume scaling of torodrops, a two-phase flow, two-dimensional (2D) axisymmetric model was developed based on our previously reported numerical frameworks.^29,30^ The model considered the inner radius of the hydrogel ring (*r*), and the thicknesses of the aqueous phase surrounding the hydrogel ring (*t*_x_ and *t*_y_), respectively (**Figure S1a** and **Note S1**). Because excess liquid was removed during compartment formation, thin aqueous layers were assumed (*t*_x_ < 100 μm and *t*_y_ < 60 μm) (**Table S2**), while *r* and *D* were varied from 100 μm to 250 μm and 200 μm to 800 μm, respectively (Figure S1b). Simulation results revealed that hydrogel ring gap was the dominant factor governing torodrop isolation (Figure 3b). When *D* was below 600 μm, the residual aqueous solution remained trapped between adjacent rings, resulting in connected drops. Increasing *D* to 600 μm enabled the formation of isolated torodrops, where variations in *r* had little impact on torodrops isolation. To validate the simulation, the hydrogel ring geometry was experimentally tuned by modulating the photomask design and the flow rate ratio of PEG and PEGDA (*Q*_PEG_:*Q*_PEGDA_) (**Figure S2**). Consistent with the simulation, increasing *D* to 600 μm produced predominantly discrete torodrops, although occasional pinned drops were observed because of fabrication imperfections. Uniform torodrops were obtained when the *D* exceeded 600 μm (Figure 3c). To ensure robust and reproducible torodrop generation, a distance (*D*) of 1 mm was used in all subsequent experiments.

The effect of hydrogel geometry on torodrop volume was further investigated through numerical simulations and experiments. As expected, simulation results revealed that the pinned drops retained larger volumes than isolated torodrops (Figure S1c). Crucially, both the experimental and simulated torodrop volumes decreased linearly with increasing *r* (R^2^ = 0.9995 and 0.9754, respectively) (Figure 3d and **Note S2**). Experimentally, increasing the flow rate ratio *Q*_PEG_:*Q*_PEGDA_ from 1:1 to 6:1 increased the *r* from 157.8 μm to 239.9 μm (**Table S3**), leading to a corresponding reduction in torodrop volume (*V*_t_) from 47.4 nL to 15.8 nL. Similarly, the simulated *V*_t_ decreased from 57.0 nL to 28.1 nL as *r* increased from 150 μm to 250 μm (**Table S4**). Although the experimental and simulated results exhibited same linear trends, the absolute experimental volumes were smaller than the simulated values, particularly at larger inner radii. This discrepancy is likely attributable to the simplified two-dimensional model and the assumptions used to estimate torodrop volume experimentally.

The uniformity and volume of the torodrops are strongly influenced by the geometry of the hydrogel rings. We next evaluated fabrication consistency. The hydrogel rings exhibited excellent uniformity across different flow rate ratios, with the normalized cross-sectional area (top view) ranging from 0.86 to 1 (Figure 3e). The slight increase in area is attributed to the progressively thinner PEGDA layer surrounding the PEG core at higher PEG flow rates, which is more susceptible to lateral photopolymerization under identical UV exposure conditions. Specifically, when the flow rate ratio *Q*_PEG_:*Q*_PEGDA_ = 1:1, the hydrogel ring width (*w*) was 279.8 μm with a coefficient of variation (CV) of 5.0%. As the flow rate ratio increased to 6:1, *w* increased to 326.7 μm with a CV of 5.7%. In all cases, the measured widths exceeded the 250 μm slit length of the photomask owing to the divergence of the uncollimated UV light during photopolymerization. The normalized fluorescence intensity of the hydrogel rings decreased linearly by 39.6% as *Q*_PEG_:*Q*_PEGDA_ increased from 1:1 to 6:1, confirming that hydrogel thickness could be controlled by tuning the flow conditions. The geometry of the LabCap system was also highly reproducible between fabrication batches. When *Q*_PEG_:*Q*_PEGDA_ = 2:1, the variation in hydrogel ring area and thickness among six independent capillaries was only 2.9% and 2.8%, respectively (Figure 3f).

The stability of torodrops was further evaluated in the absence of an oil phase, which is commonly used to stabilize the segmented droplets. A capillary fabricated at *Q*_PEG_:*Q*_PEGDA_=2:1 was continuously imaged every 15 min for 1 h following torodrops formation (**Figure S3a**). Aqueous fluorescein solution partitioned into the PEGDA hydrogel network was used to monitor torodrops stability. After torodrops formation, the fluorescence intensity along the capillary was reasonably uniform, with a CV of 7.6%. Over time, a noticeable decrease in both volume and fluorescence intensity was observed for the two hydrogel rings located at the open ends of the capillary (Figure S3b). In contrast, the hydrogel rings located in the central region maintained nearly constant fluorescence intensity and morphology throughout the observation period. After excluding the terminal rings, the fluorescence intensity of the remaining hydrogel rings remained stable after 1 h, with a CV of 7.3% (Figure 3g) and an average decrease in fluorescent intensity of 1.9%. These results demonstrated that the LabCap system provides uniform, stable, and reproducible torodrops, with evaporation effects confined to the capillary ends. Here, we used LabCaps fabricated with a *Q*_PEG_:*Q*_PEGDA_ = 2:1 for the following experiments.

### 2.4 Affinity assay

To enable affinity capture, hydrogel rings were functionalized with biotin-PEGDA. We loaded streptavidin-Alexa Fluor555 (SA-AF555) into LabCap via capillary action. After incubating for 30 min, the unbound SA-AF555 was removed from the capillary by using tissue paper, followed by multiple washing steps (**Figure 4a**). To determine the spatial distribution of binding within the annular hydrogel structure, fluorescence intensity profiles were analyzed along the x- and y-axes. Fluorescence was localized primarily at the lumen-facing surfaces of the hydrogel rings, while the hydrogel bulk and the outer surface attached to the capillary wall exhibited only background signal (**Figure S4a**). Along the x-axis, two sharp intensity peaks were observed at the edges of the ring, coinciding with the surfaces directly exposed to the SA-AF555 solution (Figure S4b). Similar but weaker peaks were observed along the y-axis. These results indicate that streptavidin-biotin binding occurs predominantly at the hydrogel-solution interface facing the lumen rather than throughout the hydrogel volume. The confinement of fluorescence to a thin boundary layer suggests a diffusion-limited binding process, where restricted transport within the polymer network limits target penetration into the hydrogel interior.^10,31^ The binding performance of the biotinylated hydrogel rings was evaluated using SA-AF555 concentrations ranging from 10 nM to 1000 nM. Fluorescence intensity increased with target concentration and reached a plateau at approximately 300 nM (Figure 4b and Figure S4c-d). A slight concentration-dependent increase was also observed in non-biotinylated control rings because of nonspecific adsorption. Nevertheless, the control signal remained significantly lower than that of the biotinylated hydrogel rings. The signal-to-noise ratio (SNR) increased with SA-AF555 concentration and reached a maximum at 300 nM, after which it decreased because of increased nonspecific fluorescence at higher SA-AF555 (Figure 4b). The washing protocol was subsequently optimized by varying the number of wash cycles. The background signal of the capillary decreased substantially between 3 and 6 wash cycles and reached a stable baseline after 10 wash cycles, with no further change in either hydrogel ring or background fluorescence (Figure 4c). In the following experiment, 15 wash cycles were performed to ensure that excess target molecules were completely removed. These results demonstrate the high specificity and effective target capture performance of the LabCap platform.

**Figure 4.**
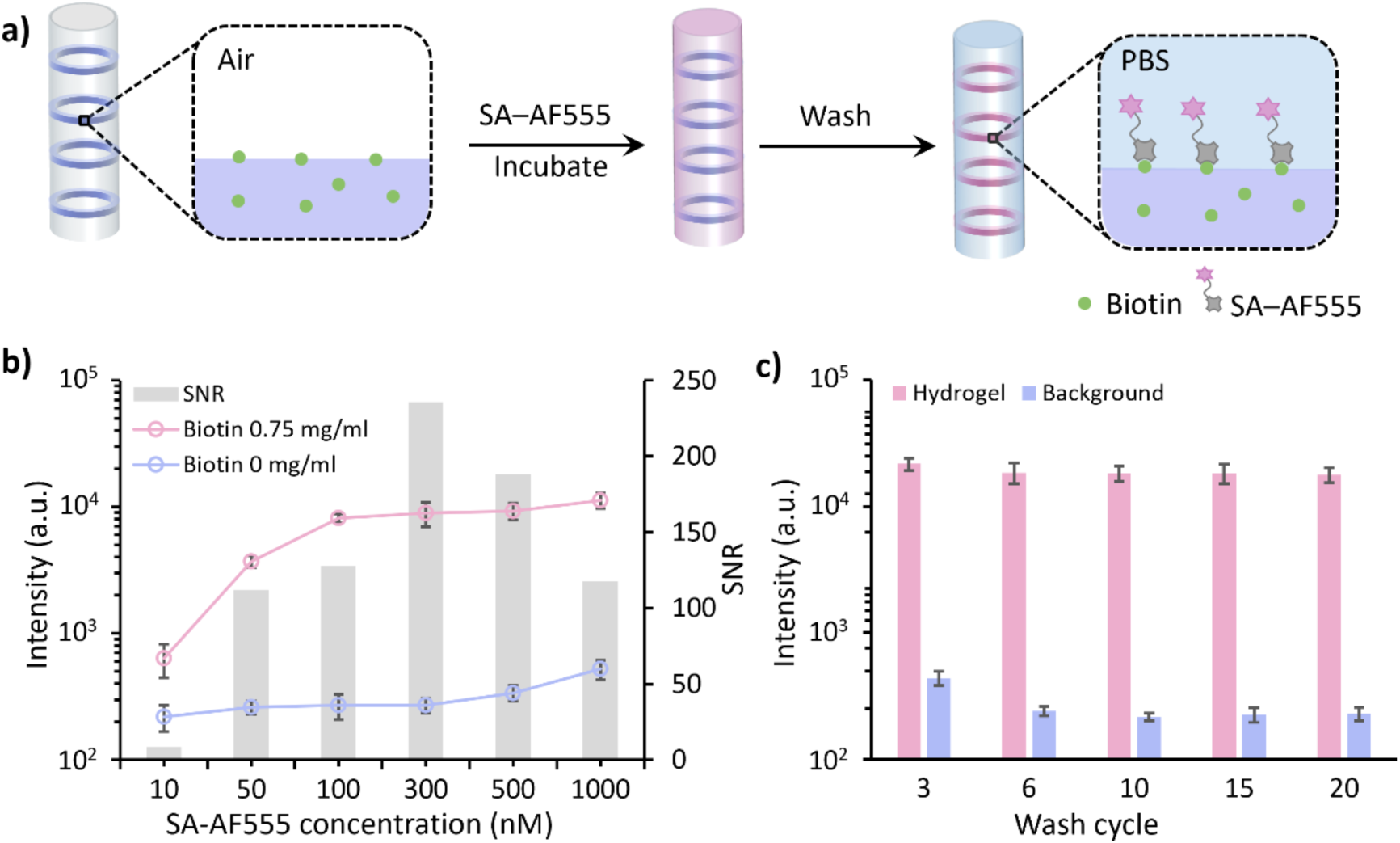
Evaluation of affinity binding on LabCap platform. (a) Schematic illustration of fluorescent streptavidin (SA-AF555) binding to biotinylated hydrogel rings, followed by washing steps. (b) Fluorescence intensity of biotinylated hydrogel rings and non-biotinylated hydrogel rings, and signal to noise (SNR) as a function of SA-AF555 concentration. (c) Effect of wash cycles on the fluorescence intensity of biotinylated hydrogel rings and background at SA-AF555 concentration of 300 nM. Error bars represent the standard deviation of fluorescence intensities measured from multiple hydrogel rings within a single capillary (n = 26).

We have investigated three incubation strategies, including periodic medium exchange (method A), static incubation (method B), and continuous flow injection (method C), to reduce the incubation time with minimal reagent consumption (**Figure 5a**). In the medium exchange protocol (method A), the SA-AF555 was loaded into the capillary, incubated for 1 min, removed, and replaced with a fresh solution for multiple cycles. In the static incubation protocol (method B), the solution was loaded once and allowed to equilibrate without disturbance. In contrast, the flow incubation protocol (method C) continuously perfused the capillary with fresh solution using a syringe pump. Because convective transport continuously replenishes target molecules at the hydrogel surface, flow incubation served as the reference condition (*Pe* = 1830, **Note S3**). Static incubation represents a depletion-limited regime, whereas periodic medium exchange provides an intermediate strategy that periodically renews reactants without requiring a larger sample volume (*Pe* = 6.5×10^4^).

**Figure 5.**
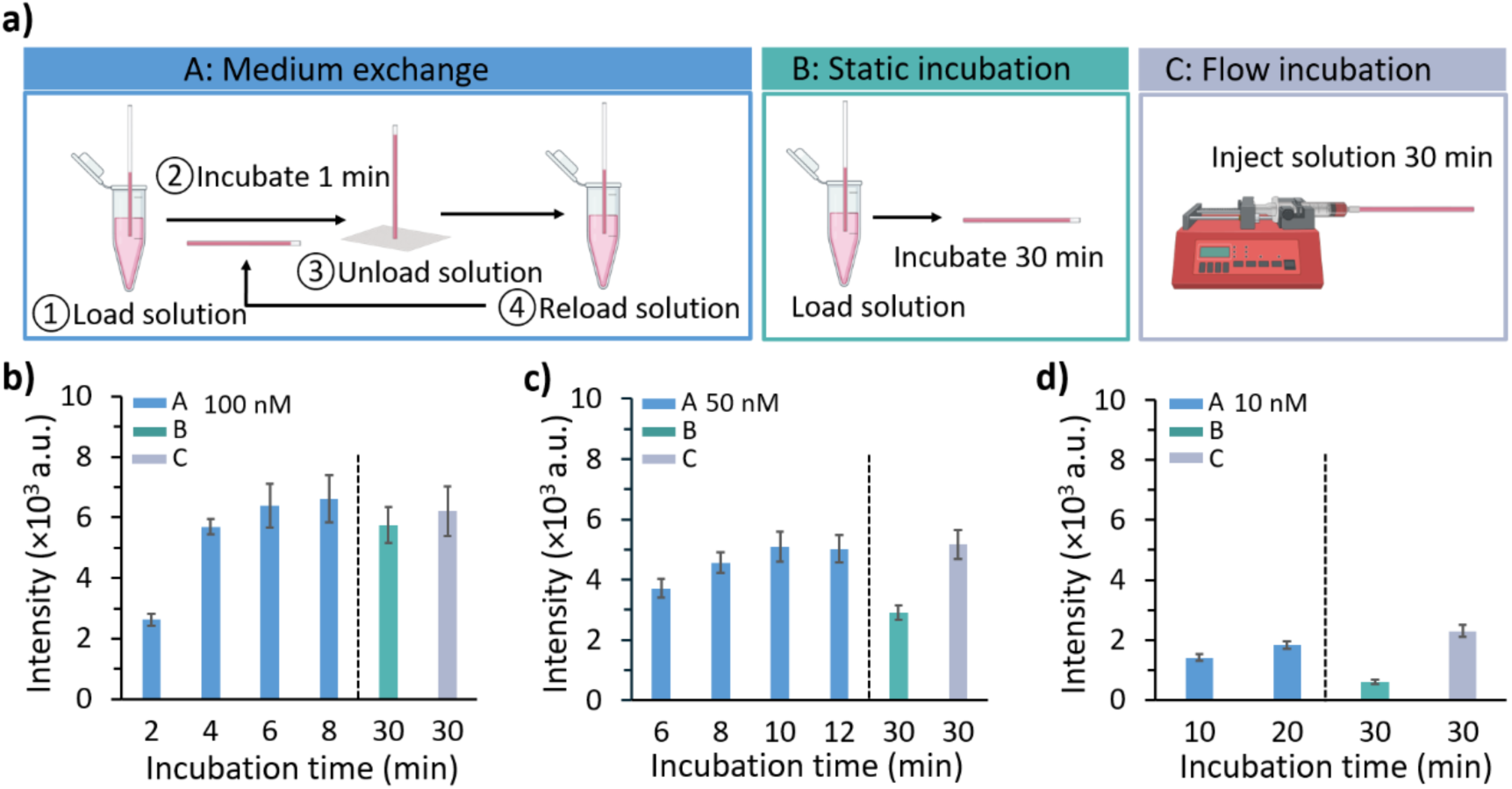
Comparison of incubation strategies for streptavidin-biotin binding in LabCap. (a) Schematic illustration of three incubation strategies: periodic medium exchange (method A), static incubation (method B), and continuous flow incubation (method C). Created with BioRender. (b-d) Fluorescence intensity of the hydrogel rings following incubation with SA-AF555 at concentrations of 100 nM (b), 50 nM (c), and 10 nM (d) using three incubation strategies. Error bars represent the standard deviation of fluorescence intensities measured from multiple hydrogel rings within a single capillary (n = 26).

We first incubated biotinylated hydrogel rings with 100 nM SA-AF555 using these three different strategies. After five medium exchange cycles (incubation time 6 min) using method A, the hydrogel rings exhibited fluorescence intensity (6.6×10³ a.u.) comparable to that observed after 30 min of flow incubation using method C (6.2×10³ a.u.) (Figure 5b). This indicated that periodic replenishment effectively accelerates binding kinetics. Notably, under the same streptavidin concentration (100 nM), 30 min of static incubation using method B produced 93% of the fluorescence intensity obtained under continuous flow, suggesting that the hydrogel surface was nearly saturated at this relatively high target concentration. More pronounced depletion-limited binding kinetics emerged at lower target concentrations. At 50 nM of SA-AF555, nine exchange cycles (incubation time 10 min) achieved 98.5% of the fluorescence intensity obtained after 30 min of flow incubation, whereas static incubation yielded only ∼56.4% of the corresponding signal (Figure 5c). When the concentration was further reduced to 10 nM, even nineteen exchange cycles (incubation time 20 min) achieved 79.9% of the fluorescence intensity obtained under flow incubation signal, while static incubation produced only 25.9% of the fluorescence intensity after 30 min (Figure 5d). The concentration-dependent behavior can be well interpreted through Fick’s first law of diffusion^32^, 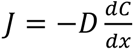, where *J* is the diffusive flux, 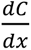 is the local concentration gradient, *D* is the diffusion coefficient. At lower target concentrations, depletion near the hydrogel surface reduces the gradient, which lowers the flux. Consequently, molecular transport slows down, requiring a longer incubation time and more medium exchange cycles to achieve a comparable signal intensity. These results demonstrate that the incubation method plays a critical role in determining binding efficiency within confined hydrogel geometries. Although continuous flow provides the most favorable mass transport conditions, it requires substantially larger reagent volumes (>300 µL per incubation step) and a syringe pump. In contrast, periodic medium exchange substantially improves molecular transport while consuming less sample volumes (96 µL for 7 cycles) in an instrument-free workflow. At relatively high target concentrations, periodic medium exchange achieves binding performance comparable to continuous flow within a short incubation time. As the target concentration decreases, additional medium exchange cycles are required to compensate for the reduced diffusive flux and maintain efficient target capture, whereas the static incubation method fails to keep up. These results demonstrate that tailoring the incubation strategy regulates mass transport and binding kinetics.

### 2.5 Enzymatic amplification assay

We next evaluated the signal amplification capability of LabCap platform using enzymatic amplification assays. Biotinylated hydrogel rings were first incubated with streptavidin-enzyme conjugates (SA-enzyme) using the optimized periodic medium exchange protocol for 10 min (nine exchange cycles), enabling efficient enzyme immobilization within the hydrogel rings. Following a washing step, the corresponding fluorogenic substrate solution was introduced into the capillary. The capillary was then inverted and the bulk solution removed, resulting in the formation of isolated torodrops around each hydrogel ring (**Figure 6a**). Two widely used ELISA enzyme systems were tested, including horseradish peroxidase (HRP) and β-galactosidase (β-gal). As illustrated in Figure 6b, HRP catalyzes the oxidation of 10-acetyl-3,7-dihydroxyphenoxazine (ADHP) into fluorescent resorufin in the presence of hydrogen peroxide, while β-gal hydrolyzes fluorescein-di-β-glucopyranoside (FDG) into fluorescein.

**Figure 6.**
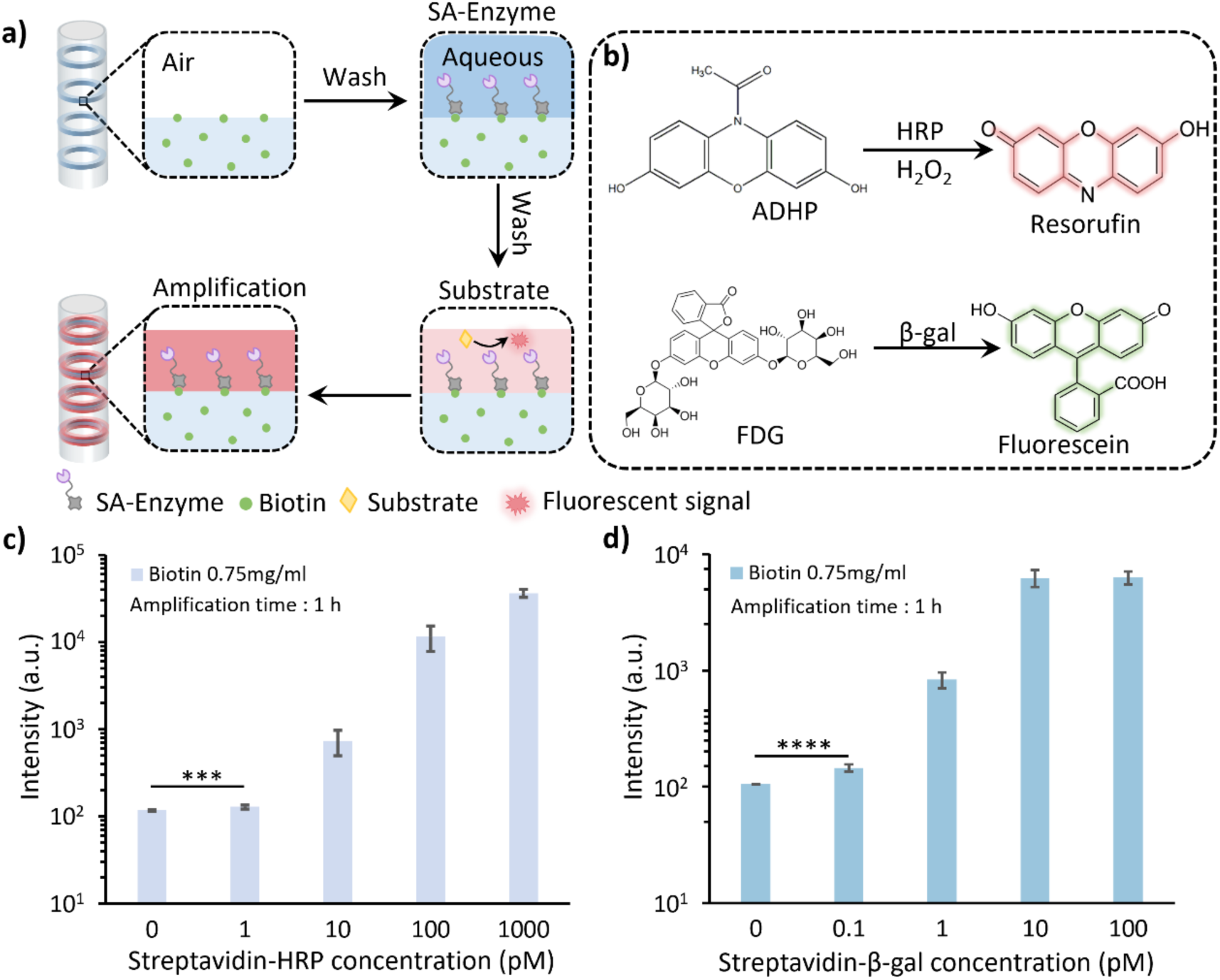
Enzymatic amplification performance of LabCap platform. (a) Schematic illustration of the enzymatic amplification assay workflow. (b) Reaction schemes of the two enzymatic amplification systems. (c, d) Mean fluorescence intensity of hydrogel rings following enzymatic amplification as a function of SA-HRP concentration (c) and SA-β-gal concentration (d). Error bars represent the standard deviation of fluorescence intensities measured from multiple hydrogel rings within a single capillary (20 ≤ n ≤ 26). *p*-values were determined by two-tailed Student t-tests. *** *p* < 0.001, **** *p* < 0.0001.

Following torodrops formation, the enzymatic amplification was initiated, resulting in the accumulation of fluorescent products within confined aqueous compartments and their partitioning into the hydrogel rings. Overlay images of fluorescence and bright-field microscopy after 1 h of amplification clearly showed localized signal generation within the hydrogel regions (**Figure S5a, b**). Quantitative analysis was performed using the mean fluorescence intensity of the hydrogel ring area. In the HRP/ADHP system, signal intensity increased with SA-HRP over the range of 1-1000 pM as the fluorescent signal accumulated over time (Figure 6c and Figure S5c). A similar concentration-dependent response was observed for β-gal/FDG system over the range of 0.1-100 pM (Figure 6d and Figure S5d). To evaluate nonspecific signal generation, non-biotinylated hydrogel rings were used as negative controls. These control rings exhibited only weak fluorescence throughout the investigated concentration range. Although the background fluorescence increased slightly, it remained lower than that of the biotinylated hydrogel rings, indicating minimal nonspecific adsorption (Figure S5e, f). The SNR increased gradually with increasing concentrations and reached a maximum at approximately 10 pM for both streptavidin-conjugated enzymes, after which it decreased because of increased nonspecific fluorescence at higher concentration. Thus, LabCap enables sensitive detection at picomolar concentrations while maintaining a dynamic range spanning three orders of magnitude across both enzymatic amplification systems.

### 2.6 Detection of biomarkers

As a proof of concept, the LabCap platform was applied to detect CRP and NT-proBNP using a sandwich ELISA format (**Figure 7**). In previous affinity assays, the medium was exchanged every one minute to enhance mass transport. However, since the ELISA process requires multiple incubation and washing steps, the exchange interval was extended to two minutes to ensure smoother operation while maintaining a total incubation time of sixteen minutes (seven exchange cycles) (Figure 7a). During the incubation process, sandwich immunocomplex was assembled on the biotinylated hydrogel ring. Removal of the excess ADHP substrate solution generated torodrops around each hydrogel rings. The HRP confined the torodrops converted ADHP into resorufin to amplify fluorescence signals for biomarker quantification (Figure 7b).

**Figure 7.**
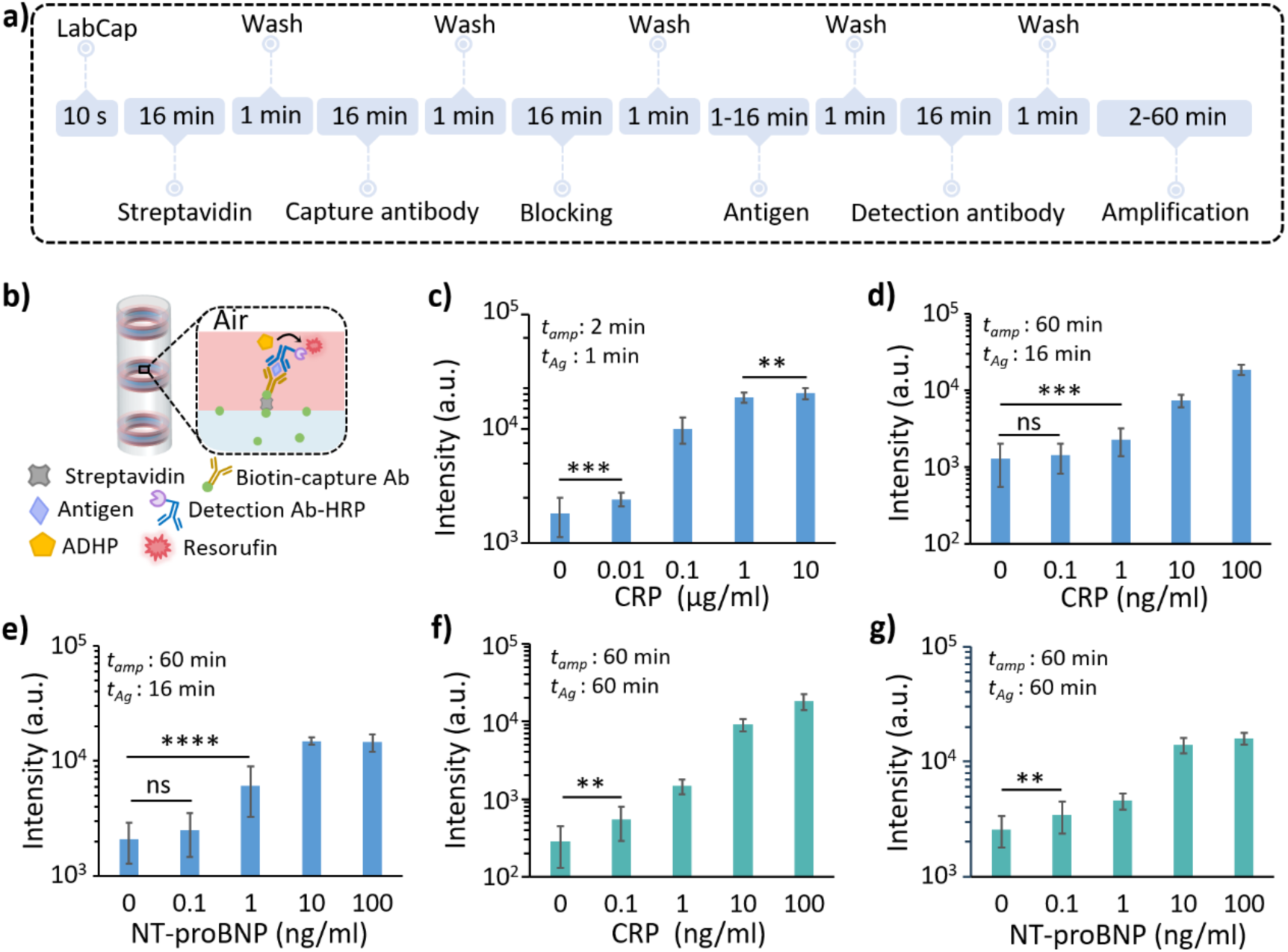
Biomarkers detection using LabCap platform. (a) Schematic illustration of sandwich ELISA workflow employing medium exchange protocol. (b) Schematic illustration of sandwich immunocomplex formation within the hydrogel rings. (c-e) Mean fluorescence intensity of hydrogel rings following enzymatic amplification as a function of CRP concentration (c, d) and NT-proBNP concentration (e) using medium exchange protocol. (f, g) Mean fluorescence intensity of hydrogel rings following enzymatic amplification as a function of CRP concentration (f) and NT-proBNP concentration (g) using static incubation protocol. Error bars represent the standard deviation of fluorescence intensities measured from multiple hydrogel rings within a single capillary (20 ≤ n ≤ 26). *p*-values were determined by two-tailed Student t-tests. ns: *p* ˃ 0.05, ** *p* < 0.01, *** *p* < 0.001, **** *p* < 0.0001. *t_amp_* and *t_Ag_* denote the amplification time and the antigen incubation time, respectively.

To demonstrate clinical applicability, CRP detection was first evaluated across a wide concentration range (0.01-10 µg/mL) using medium exchange protocol (method A), encompassing both normal (<1 µg/mL) and elevated levels associated with inflammatory conditions.^33,34^ As shown in **Figure S6a**, the fluorescence intensity from hydrogel rings rapidly reached a plateau within 5 min of amplification for CRP concentrations above 0.1 µg/mL. Moreover, plateau values slightly decreased with increasing CRP concentration, which may result from excessive antigen binding, which can hinder efficient sandwich complex formation. To alleviate this effect, the CRP incubation time was reduced to 1 min while keeping other assay parameters unchanged. Under this condition, signal saturation was still observed at longer amplification times; however, the intensity increased monotonically with CRP concentration (Figure S6b, c). Using an amplification time of 2 min, the LabCap system enabled reliable detection of CRP down to 0.01 µg/mL while maintaining quantitative detection across the clinically relevant concentration range of 1-10 µg/mL (Figure 7c and Figure S6c).

The system capability for low-concentration detection of CRP and NT-proBNP was also examined using the medium exchange protocol (method A), which employed sixteen minutes of incubation steps and 60 min amplification. Under these conditions, both biomarkers could be detected at low ng/mL levels. However, when the biomarker concentration decreased below 0.1 ng/mL, the measured signals were no longer statistically distinguishable from blank control (Figures 7d, e). These results indicate that although the medium exchange protocol enables fast analysis, the short incubation periods are not optimal for detecting biomarkers at very low concentrations. To further explore the versatility of LabCap, a static incubation protocol (method B) employing 1 h incubation steps for biomolecular binding was implemented. Under these conditions, the sensitivity of assay was improved, enabling detection of both CRP and NT-proBNP down to 0.1 ng/mL (Figure 7f, g). The enhanced performance is attributed to the longer incubation period, which provides sufficient time for antigen and antibody interaction to approach equilibrium and increase target capture efficiency. These results demonstrate that LabCap can be readily adapted to different analytical requirements by adjusting the incubation conditions. The medium exchange protocols enable accelerated detection of high-abundance biomarkers, whereas the static incubation protocol provides enhanced sensitivity for low-abundance targets.

In addition to sensitivity and assay time, reagent consumption is an important practical consideration for diagnostic applications (**Table S5**). Notably, the static incubation protocol (method B) required only 12 μL of reagent per assay step, maintaining sensitive detection of low-concentration biomarkers. The medium exchange protocol (method A) consumed a total of 96 µL of reagent per assay step. Besides, the wash buffer volume was dramatically reduced to 1 mL regardless of the incubation protocols.

The storage stability of LabCap was also evaluated. The hydrogel rings were first functionalized with CRP capture antibodies and subsequently stored in 1% BSA solution at 4 °C for one week. Following storage, the pre-functionalized LabCap was directly used for CRP detection over the clinically relevant concentration range using the medium exchange protocol (method A) (**Figure 8a**). The fluorescence intensity distribution shifted to the right with increasing CRP concentration, indicating that the concentration-dependent response of the assay was preserved after storage (Figure 8b). Consistently, the mean fluorescence intensity, averaged across three parallel experiments, increased with increasing CRP concentration (Figure 8c). Although the stored LabCap exhibited a slight reduction in fluorescence intensity compared with freshly prepared devices, it retained approximately 88.1% of the original fluorescence response at 1 µg/mL. These results demonstrate that LabCap maintained its functional performance after at least one week of storage. Furthermore, because the hydrogel rings can be pre-functionalized prior to use, the assay operation time for the end user is reduced from over 4 h (for the complete workflow) to less than 30 min using the pre-functionalized LabCap.

**Figure 8.**
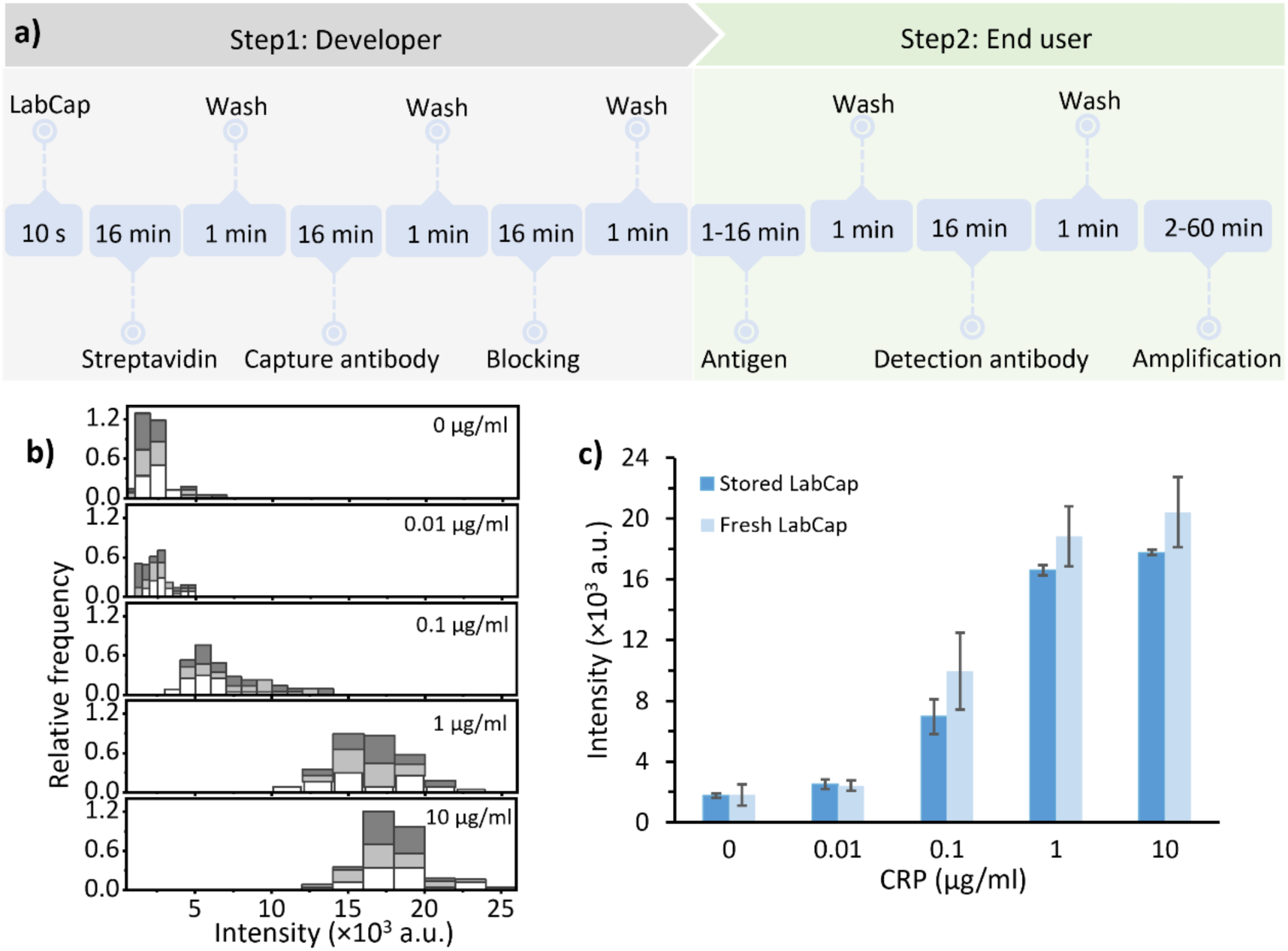
Stability evaluation of LabCap platform. (a) Schematic illustration of the storage stability test using medium exchange protocol. (b) Fluorescence intensity distribution of hydrogel rings following CRP detection after storage in 1% BSA at 4 °C for one week. Stacked histograms represent pooled data from three independent experiments, with each color corresponding to a different experiment. (c) Comparison of the mean fluorescence intensity of hydrogel rings as a function of CRP concentration between stored LabCap and fresh LabCap. The stored LabCap data was obtained from three independent capillaries. For each capillary, the mean fluorescence intensity was calculated from 22 hydrogel rings, and data are presented as mean ± standard deviation (N = 3). The fresh LabCap data are reproduced from Figure 7c for comparison. For the fresh LabCap, the mean intensity was calculated from multiple hydrogel rings within a single capillary (20 ≤ n ≤ 26), and error bars represent the standard deviation of fluorescence intensities measured from those hydrogel rings.

## 3. Conclusion

In summary, we developed a LabCap platform comprising functionalized hydrogel rings fabricated using a reconfigurable stop-flow lithography system. LabCap enables the spontaneous formation of nanoliter-scale reaction compartments following the loading and unloading of aqueous solutions, eliminating the need for external instruments or an immiscible oil phase. These isolated compartments provide a simple and efficient environment for biomolecular assays. Using LabCap, we systematically investigated the influence of incubation strategies on biomolecular interactions. The periodic medium exchange effectively alleviates local depletion and enhances mass transport, enabling detection of CRP and NT-proBNP at concentrations as low as 1 ng/mL within short time. In contrast, static incubation provided enhanced target capture efficiency and enabled detection down to 0.1 ng/mL. These results demonstrate that assay speed and sensitivity can be tuned by adjusting the incubation protocol to meet different analytical requirements. In addition, LabCap offers several practical advantages, including low fabrication cost (< €1 per device), low reagent consumption (< 100 μL per assay step), minimal wash buffer usage (≈ 1 mL) and stable performance after 1 week storage. These features make LabCap a versatile and accessible platform for quantitative biomarker detection.

## 4. Experimental section

### 4.1 3D printing of microfluidic device and photomask

The microfluidic device and photomask were designed using SolidWorks software (Student Edition). The overall length and width of the microfluidic device were 16.5 mm and 5 mm (**Figure S7a**). The inlet ports have a tapered diameter of 1.65 mm, which is reduced to 1.4 mm for the tight insertion of polytetrafluoroethylene (PTFE) tubing (ID: 1 mm, OD: 1.59 mm, TECHLAV). The inner and outer channel widths were 0.8 mm and 0.6 mm. The wall thickness between the two channels was 0.5 mm. The diameter of the outer port was 0.7 mm. The outlet of the 3D-printed device incorporated a recessed region designed to accommodate a short segment of silicone tubing, which provided a compliant and detachable interface to the glass capillary. This interface allowed the fully developed coaxial flow to enter the capillary while enabling facile removal and replacement of the capillary between fabrication runs. The recessed area has a tapered diameter of 2.2 mm, which is reduced to 1.8 mm for the connection of a glass capillary. The overall height, length, and width of the photomask were 5 mm, 43.80 mm and 13.80 mm. The length and gap of the slit were 250 μm and 1 mm. (Figure S7b). The 3D printing process has been described extensively in previous publications.^35^ A 3D printer (Phrozen Aqua Mini 8K, Phrozen Tech Co., Ltd.) was used in this study. The microfluidic device was printed using transparent resin (Anycubic Clear Resin) and the photomask was printed using Aqua grey resin (Phrozen). Briefly, the designed stereolithographic format file was imported into Slice software (CHITUBOX Basic). To obtain a fully closed 3D device and photomask, the layer thickness and exposure time were adjusted. Specifically, 40 μm layer thickness and an exposure time 2.5 s were set for the device printing. For the photomask printing, the layer thickness and exposure time were 50 μm and 0.95 s. After printing, the devices and photomask were cleaned with isopropyl alcohol (VWR, Germany), and dried with nitrogen. After 5 min post-exposure in UV bath, they were ready to use.

### 4.2 Development of LabCap

#### 4.2.1 Surface silanization of glass capillary

To ensure the hydrogel rings adhere firmly to the inner surface of glass capillary (World Precision Instrument, 1B100-3), the glass capillary was silanized by 3-trimethoxysilyl propyl methacrylate (TMSPMA) (440159, Sigma-Aldrich). The glass capillary was soaked in 0.1 M NaOH (43617, Sigma-Aldrich) solution and sonicated (Ultrasonic bath, VWR) for 30 min. After rinsing with DI water and drying with N_2_, the glass capillary was soaked into a solution comprising 2 : 3 : 5 (v/v/v) TMSPMA: acetic acid (695092, Sigma-Aldrich) : DI water for 1 h.^36^ Then the glass capillary was rinsed with DI water and acetone (179124, Sigma-Aldrich). Lastly, the glass capillary was dried in an oven 110 °C for 30 min to facilitate covalent siloxane bond formation on the surface of the glass capillary. After silanization modification, the glass capillary connected with 3D printed microfluidic device for the establishment of LabCap by a short silicon tubing (ID: 0.8 mm, OD: 1.9 mm, Amazon).

#### 4.2.2 Experiment setup

The PTFE tubing was glued to microfluidic device two inlet ports. Another side of tubing was connected to filled syringe using flangeless fitting and union connectors. A holder was printed to mount the microfluidic device, 3D printed photomask, and align glass capillary. The holder was directly fixed on the surface of UV light (Omnicure AC450, Excelitas Technologies) using screw. The UV light passes through the slits of the photomask to selectively expose on the streams within glass capillary. The syringe pump (Chemyx) and UV light source are connected to the I/O-B module and controlled by custom LabVIEW (Figure 2c).

#### 4.2.3 Stop-flow lithography for hydrogel rings fabrication

Typically, a curable phase consists of 60% w/w PEGDA (Mw 575, 437441, Sigma-Aldrich), 40% w/w DI water. The inert phase consists of 60% PEG (Mw 200, P3015, Sigma-Aldrich), 40% DI water. The photoinitiator (2-hydroxy-2-methylpropiophenone, Darocur 1173, 405655, Sigma-Aldrich) was added to these two phases at 5% (v/v). The PEGDA phase and PEG phase were co-injected into the outer channel and inner channel of device by syringe pump at total flow rate 200 μL/min. For the fabrication of fluorescent hydrogel rings, 0.006% w/w methacryloxyethyl thiocarbamoyl rhodamine B (red fluorescent dye, Polysciences) was added to the PEGDA phase. The biotinylated hydrogel rings were prepared by adding 62.5 μL of acrylate-PEG-biotin (Biopharma PEG, Mw 5k) dissolved in DMSO (100 mg in 1.66 ml, 34869, Sigma-Aldrich) into 5 ml of PEGDA phase. The final concentration of biotin-PEGDA in PEGDA phase was 0.75 mg/mL. After the co-flow stream formed, the syringe pump stopped. After a short delay time 2 s, the stream was fully stopped and exposed to UV light (exposure time: 0.2 s, voltage: 5 V) (Figure 2c). The hydrogel rings were cured and adhered on the inner surface of the glass capillary. When single cycle of stop flow lithography was done, the glass capillary was easily exchanged to new one by inserting it into the recessed area of the device. One fabrication cycle is completed within ∼5 s. After washing with 70% ethanol (1.00983, Sigma-Aldrich) and DI water, the engineered glass capillaries were stored in water at 4 °C for later experiments. The total length of the glass capillary was 75 mm. The exposure length of the area was 43.80 mm. The excess bare part was cut by a tungsten carbide pen (Amazon) along a reference ruler. The total length of the experimental LabCap was 45 mm.

Given that the hydrodynamic radius of a 5 kDa biotin-PEGDA molecule is approximately 1.1 nm, the biotin moieties are geometrically available for streptavidin binding when situated within an effective surface layer of less than 2.2 nm from the hydrogel ring’s inner boundary. Based on precursors composition (0.75 mg/mL biotin-PEGDA), we estimated the binding site density is 6.1×10^−9^ mol/m^2^ for the hydrogel rings with an inner diameter and width of 300 μm.

### 4.3 Simulation of torodrops formation

We used the “Two-Phase Flow, Phase Field” CFD module to simulate the torodrops formation within glass capillary. To optimize computing time *T_sim_*, we established a two-dimensional axisymmetric model. This model comprised a glass capillary and two independent hydrogel rings with a microscale gap. Both the capillary and hydrogel rings are hydrophilic. The contact angles of the capillary and hydrogel rings were 15° and 5°, respectively. In the initial stage of the model, all surfaces of the hydrophilic capillary and hydrogel rings were surrounded by a thin layer of aqueous solution, while the remaining space was filled with air. The interfacial tension between water and air was 0.07 N·m^−1^. The density and dynamic viscosity of the aqueous phase were 1000 kg·m^−3^ and 0.001 Pa·s, and those of the air phase were 1.2 kg·m^−3^ and 1.8×10^−5^ Pa·s, respectively. To trace the interface accurately, the water-air interface thickness was set to one-tenth of the default value in COMSOL. By adjusting the inner radius of the hydrogel rings (*r*), the thicknesses of the aqueous solution surrounding the hydrogel rings (*t*_x_ and *t*_y_), as well as the distance between adjacent hydrogel rings (*D*), we can simulate torodrops formation under different conditions.

### 4.4 Affinity binding on LabCap platform

The engineering capillary was taken out from the fridge and washed with 0.5% Pluronic F127 (P2443, Sigma-Aldrich) by capillary force. In the medium exchange method, the SA-AF555 (S32355, Thermo Fisher Scientific) was loaded into the LabCap by capillary force, incubated for 1 min, and then unloaded before introducing a fresh solution. Different exchange periods were applied according to the concentration of SA-AF555. In the static incubation method, the LabCap incubated with SA-AF555 for 30 min without interrupt after loading solution. In the flow incubation method, a syringe pump was used to inject the SA-AF555 through the capillary at 10 μL/min for 30 min, ensuring a constant supply of target molecules. After incubation, the LabCap was washed 15 cycles with 0.5% Pluronic F127. In wash test experiments, the static incubation method was implemented. After incubating 300 nM SA-AF555, the LabCap was washed with different cycles using 0.5% Pluronic F127. The signal-to-noise ratio (SNR) was calculated as (S - B) / σ_B_, where S and B are the mean fluorescence intensity of the biotinylated rings and non-biotinylated rings, respectively, and σ_B_ is the standard deviation of the non-biotinylated rings.

### 4.5 Enzymatic amplification assay on LabCap platform

The hydrogel rings were first incubated with streptavidin-enzyme conjugates (N100, S931, Thermo Fisher Scientific) using periodic medium exchange method for 10 min (9 exchange cycles), following washing step. Then the corresponding substrate was loaded and unloaded from other side of capillary. In the β-gal amplification reaction, the FDG (F1179, Thermo Fisher Scientific) solution at 100 μM in 0.1% Pluronic F127 was used as substrate. For the HRP amplification reaction, the substrate was the QuantaRed solution (15159, Thermo Fisher Scientific) prepared following the instructions of the vendor (at 50 : 50 : 1 ratio of enhancer solution, stable peroxide, and ADHP concentrate, respectively).

### 4.6 Sandwich ELISA performance on LabCap platform

We developed a sandwich ELISA for CRP and NT-proBNP detection using commercial monoclonal antibodies. The monoclonal capture antibody and monoclonal detection antibody specific to two types of protein were ordered from HyTest. The capture antibody was conjugated with biotin, and the detection antibody was conjugated with HRP, respectively, using commercial conjugation kits following vendor instructions (Abcam ab201795, ab102890). Typically, in the static incubation ELISA protocol, biotinylated hydrogel rings are incubated with 10 μg/mL streptavidin (434302, Thermo Fisher Scientific) solution for 30 min, followed by a wash step. Then, hydrogel rings incubated with 10 μg/mL biotin-conjugated capture antibody for 1 h to complete the immobilization step, where unbound antibodies were removed through washing step. Next, the hydrogel rings were incubated with protein-free blocking buffer (37572, Thermo Fisher Scientific) for 1 h followed by washing. To test the detection of biomarkers, the hydrogel rings were incubated with a specific protein with varying concentrations of human recombinant NT-proBNP or CRP for 1 h with subsequent washing. Following, the 0.5 μg/mL HRP conjugated detection antibody was incubated with hydrogel rings for 1 h. After the washing step, the QuantaRed assay solution was loaded and unloaded from another side of the capillary immediately. In the negative group, the hydrogel rings were incubated with PBS only; all the other assay steps were kept same as the positive groups. For medium exchange ELISA, the assay workflow, reagents and concentrations were identical to those used in static incubation ELISA protocol described above. The only modification was the incubation strategy, in which incubation step was performed using periodic medium exchange for 16 min (seven exchange cycles with a 2 min interval). In the stability test experiment, the hydrogel rings were conjugated with capture antibody and stored at 1% BSA solution (A9418, Sigma-Aldrich) at 4 °C. After one week of storage, the capillaries were removed from storage and subjected to the remaining sandwich ELISA steps.

### 4.7 Image characterization

Bright field and fluorescence images of LabCap were acquired using a microscope (Thunder imager, Leica Microsystem). Hydrogel ring dimensions and fluorescence intensities were quantified using ImageJ software.

## Author contributions

Y. Y. and G. D. conceived and designed the study. Y. Y. fabricated the devices, performed the experiments, and analysed the data. M. U. A. and M. A. S. contributed to 3D printing of the device, the stop-flow lithography process and fabrication of functional capillaries, and running/optimizing the bioassay. X. S., Y. H. and L. W. contributed to the numerical simulations. Y. Y. wrote the manuscript with input from all authors. G. D. supervised the project and acquired funding. All authors have read and approved the final version of the manuscript.

## Data availability

The data supporting this article have been included as part of the Supplementary Information. Additional data that support the findings of this study are available from the corresponding author upon reasonable request.

## Acknowledgements

Y. Y. acknowledges the China Scholarship Council (CSC) for a doctoral scholarship. The authors acknowledge the support by German Research Foundation (DFG, project # 504195013). Schematics were created with BioRender.

## Supporting Information

**Figure S1.**
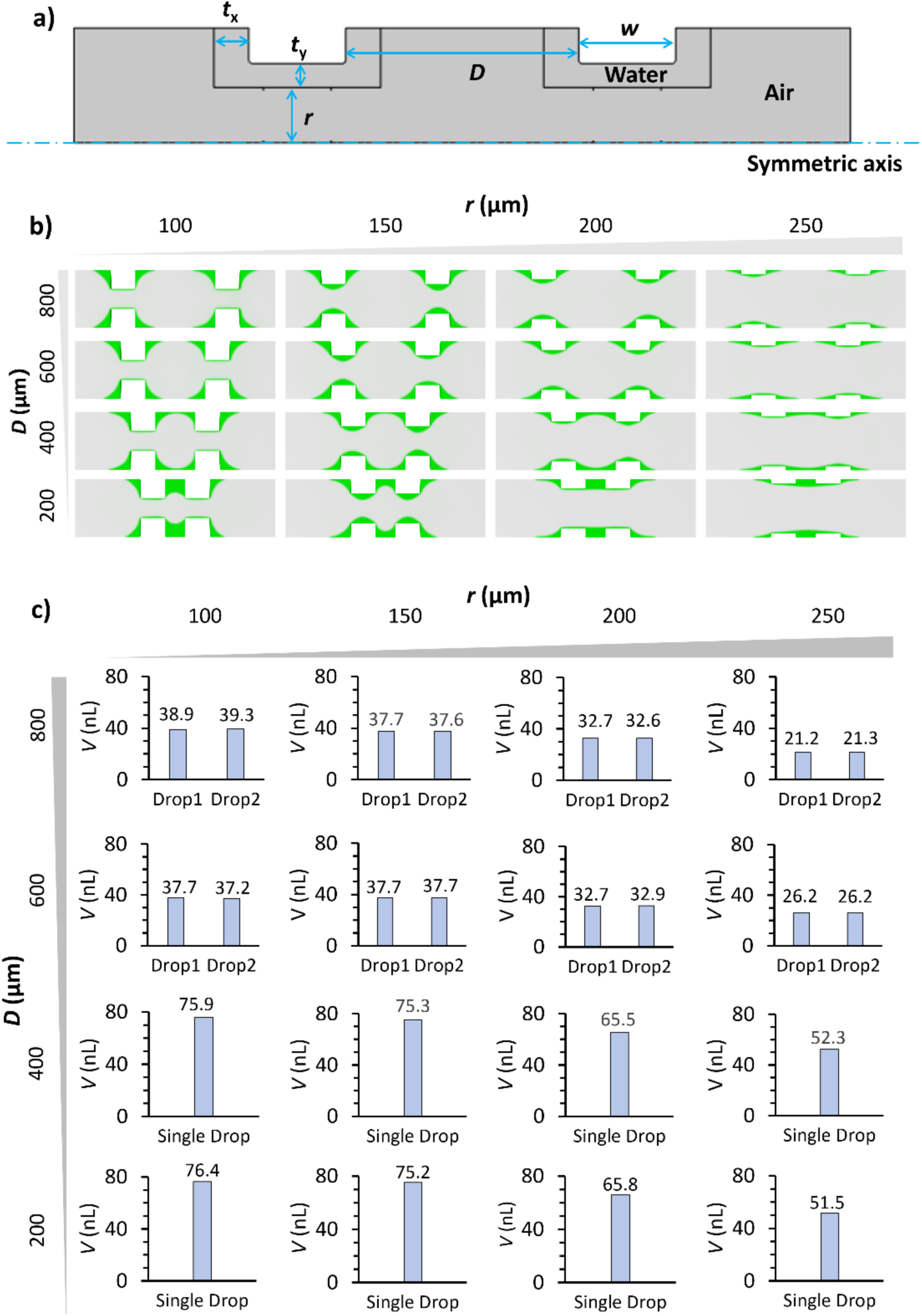
Numerical simulation of torodrop formation. (a) Schematic illustration of simulation geometric parameters used in the model. (b) Simulated torodrops formation within capillary under varying inner radius of hydrogel rings (*r*) and adjacent gap distance (*D*). (c) Torodrop volumes calculated from the simulation model (*V*_s_) excluding the volume occupied by the hydrogel rings (*V*_in_).

**Figure S2.**
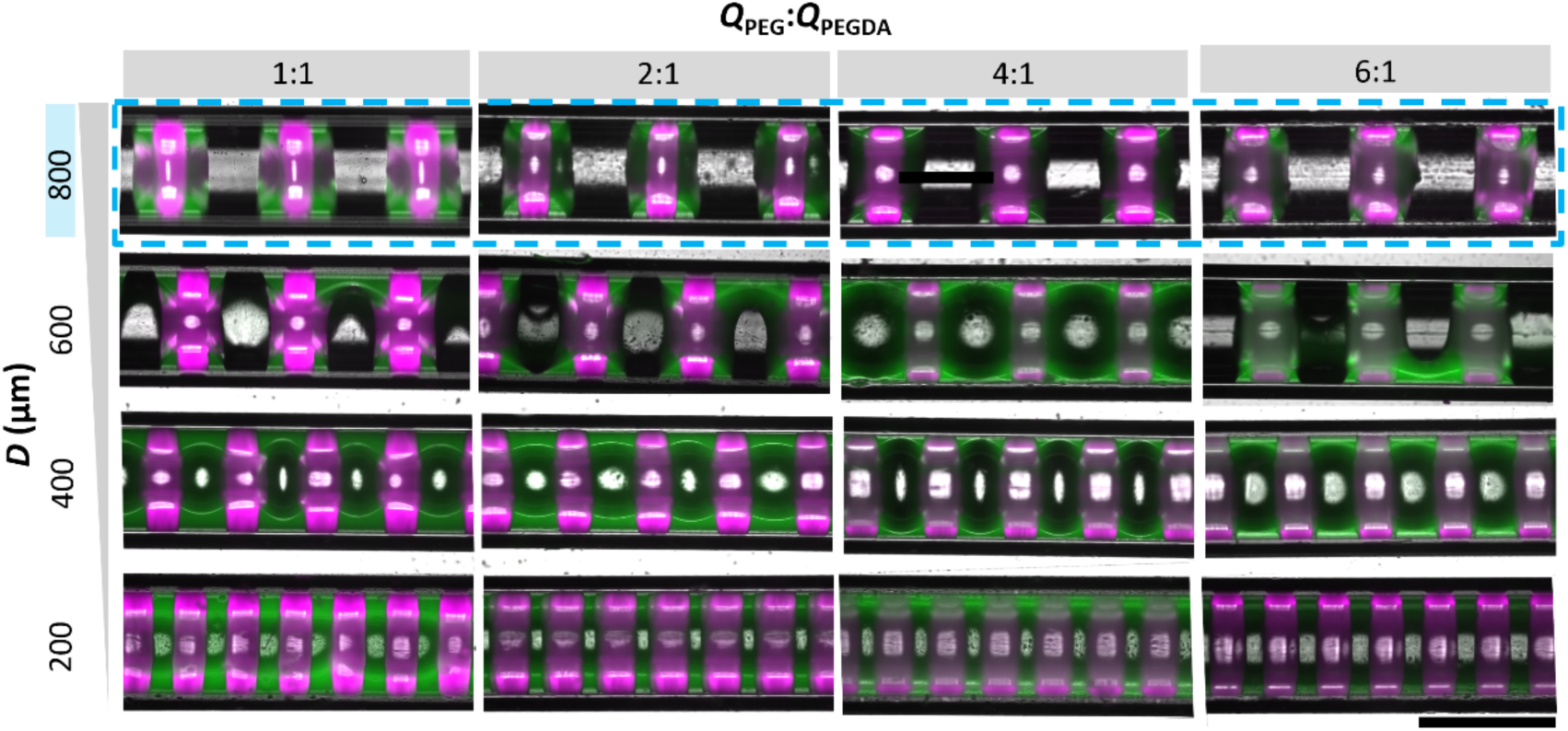
Merged images of bright-field and fluorescence channels showing torodrops formation within a capillary. The green-emitting drops were generated around red-fluorescent hydrogel rings featuring varying adjacent hydrogel distance (*D*) and inner radius. The geometric variations in the hydrogel rings were achieved using different photomask designs and core-to-shell flow rate ratios (*Q*_PEG_:*Q*_PEGDA_). The scale bar represents 1 mm.

**Figure S3.**
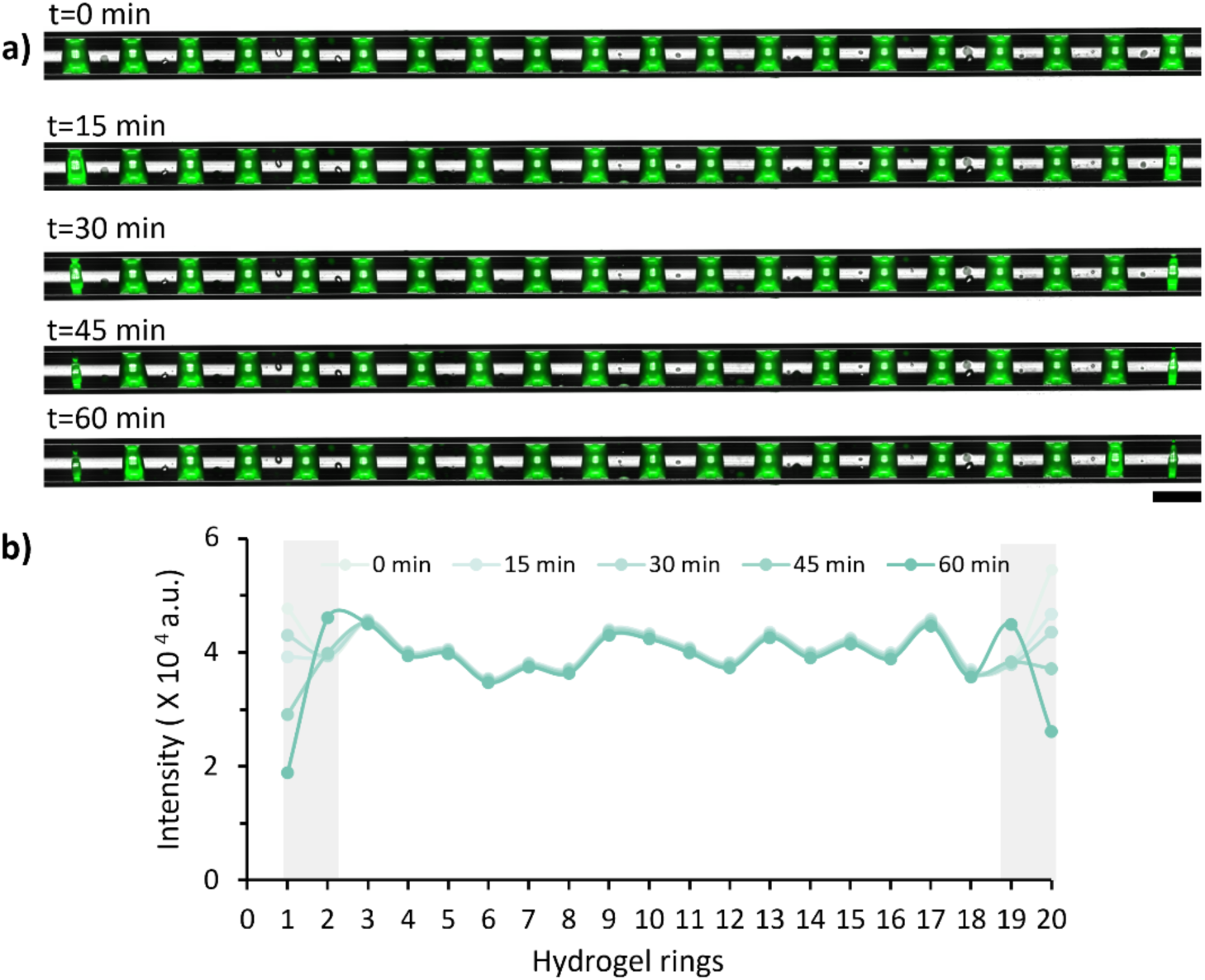
The evaporation of torodrops was evaluated at room temperature within 1 hour. (a) Merged images of bright field and fluorescence channels showing torodrops at different time points under room temperature. (b) Fluorescence intensity of hydrogel rings within capillary at different time points. The scale bar represents 1 mm.

**Figure S4.**
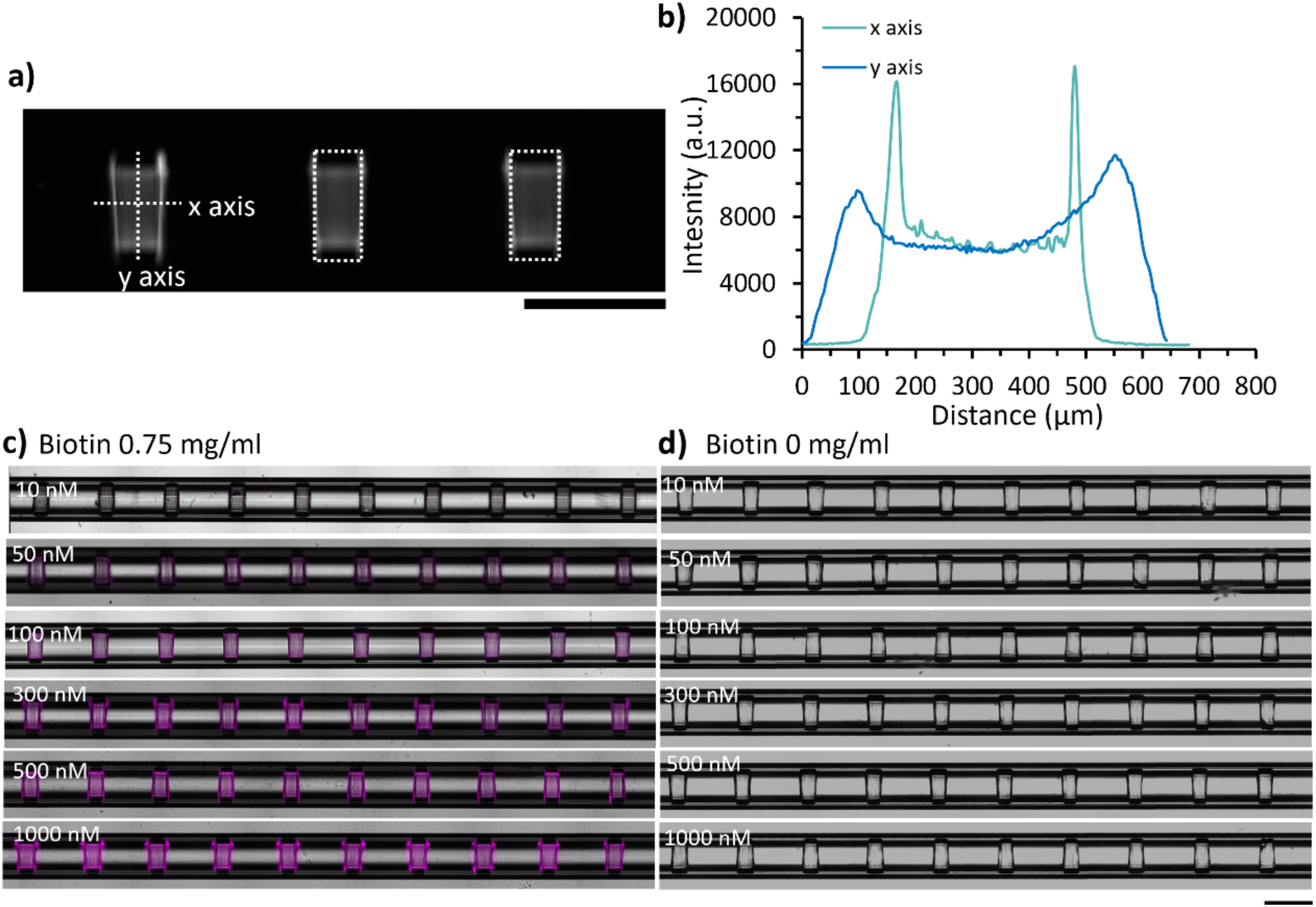
Affinity assay performance of LabCap. (a) The fluorescence image of hydrogel rings following incubation with 300 nM streptavidin-Alexa Fluor 555 (SA-AF555). (b) Fluorescence intensity profiles along the x axis and y axis of the hydrogel ring shown in panel a. (c, d) Merged images of bright field and fluorescence channels showing biotinylated hydrogel rings (c) and non-biotinylated hydrogel rings (d) following incubation with various concentrations of SA-AF555. The scale bar represents 1 mm.

**Figure S5.**
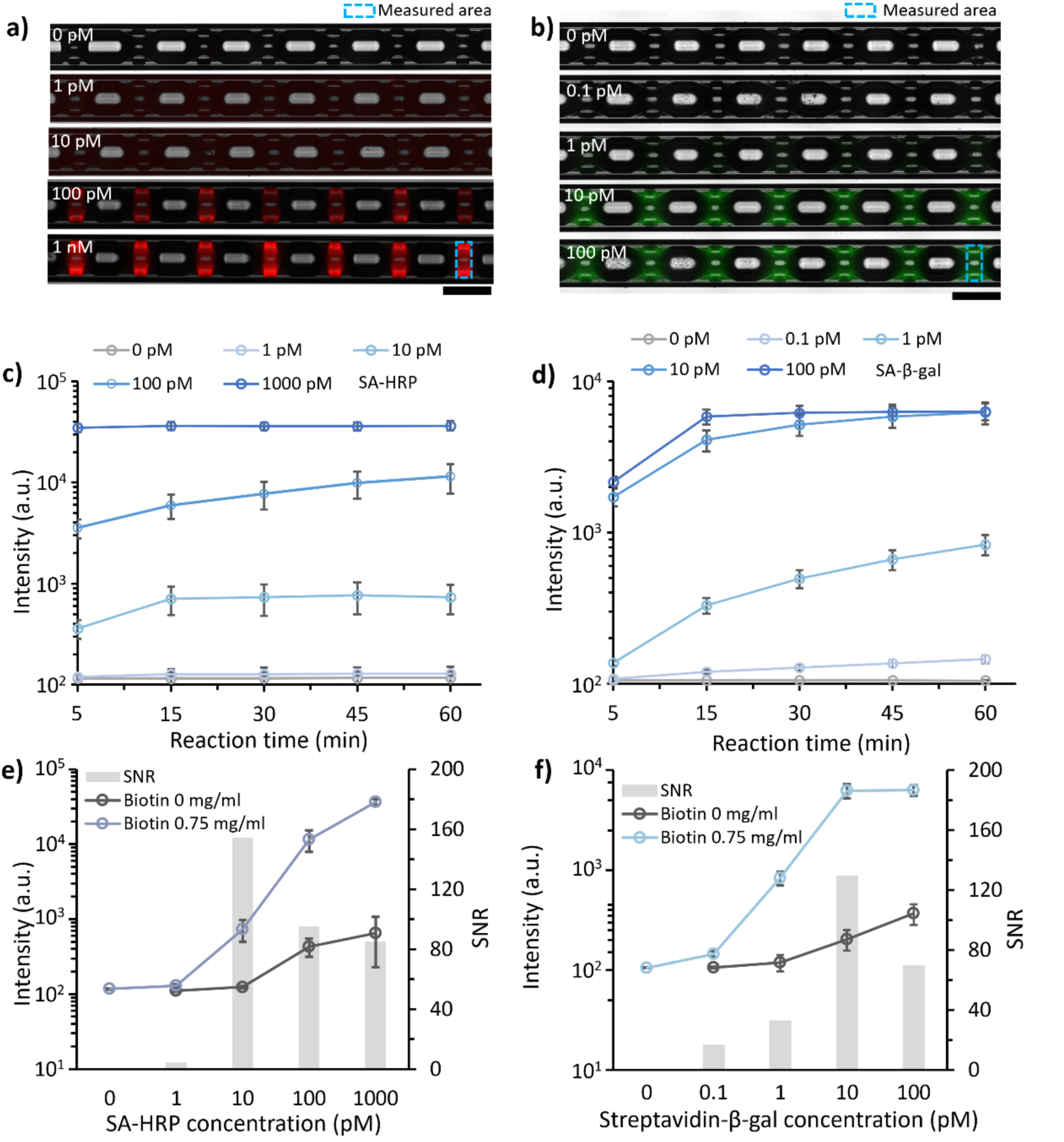
Enzymatic amplification assay performance of LabCap. Merged images of bright field and fluorescence channels showing biotinylated hydrogel rings incubating with various concentrations of SA-HRP (a) and SA-β-gal (b) after 60 min of amplification. Scale bars represent 1 mm. (c, d) Mean intensity of biotinylated hydrogel rings incubated with various concentrations of SA-HRP (c) and SA-β-gal (d) as a function of amplification time. (c) At 1000 pM SA-HRP, the fluorescence signal increased rapidly and reached a plateau within 5 min. In contrast, at lower enzyme concentrations (≤ 100 pM), signal growth was slower and continued to increase till 60 min. These results indicate that the amplification kinetics are strongly dependent on enzyme concentration. (d) At 100 pM of SA–β-gal, the fluorescence signal reached saturation within 15 min. At lower concentrations, the signal continued to increase throughout the 1 h amplification period. The signal saturation at higher concentrations, i.e., 1000 pM SA-HRP (c) and 100 pM of SA–β-gal (d), can be attributed to a complete consumption of the substrate solution within the confined torodrop volumes. (e, f) Mean fluorescence intensity of biotinylated and non-biotinylated hydrogel rings, and signal to noise (SNR) as a function of SA-HRP (e) and SA-β-gal (f) concentration after 60 min of amplification. Error bars represent the standard deviation of fluorescence intensities measured from multiple hydrogel rings within a single capillary (20 ≤ n ≤ 26).

**Figure S6.**
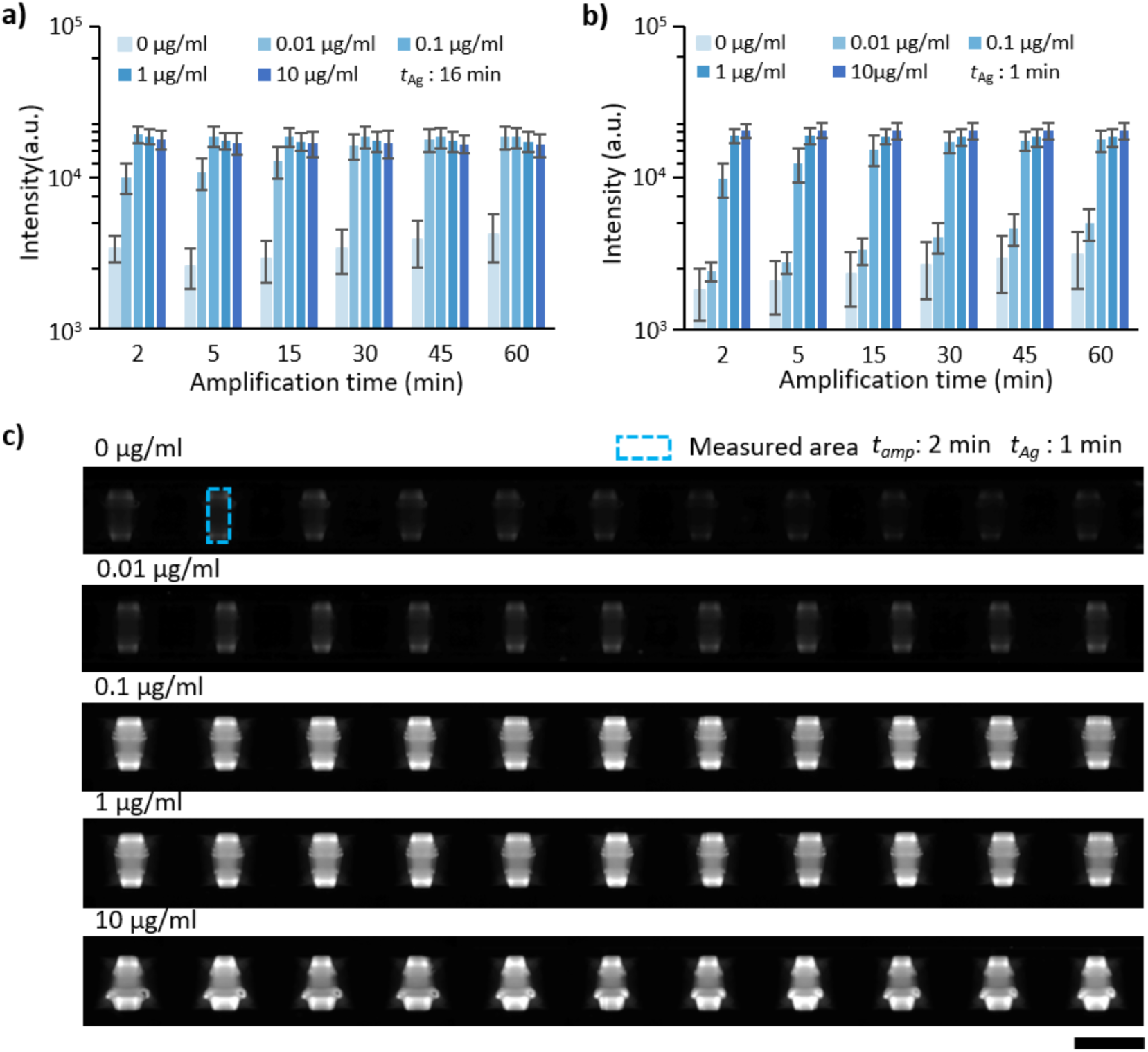
CRP detection on LabCap. (a, b) Mean intensity of hydrogel rings measured at various CRP concentrations following different amplification times. Antigen incubation time (*t*_Ag_) was fixed at 16 min (a) and 1 min (b). Error bars represent the standard deviation of fluorescence intensities measured from multiple hydrogel rings within a single capillary (20 ≤ n ≤ 26). (c).Fluorescence images showing concentration-dependent response of hydrogel ring fluorescence intensity to CRP. Amplification time (*t_amp_*) and antigen incubation time (*t_Ag_*) were 2 min and 1 min. The scale bar represents 1 mm.

**Figure S7.**
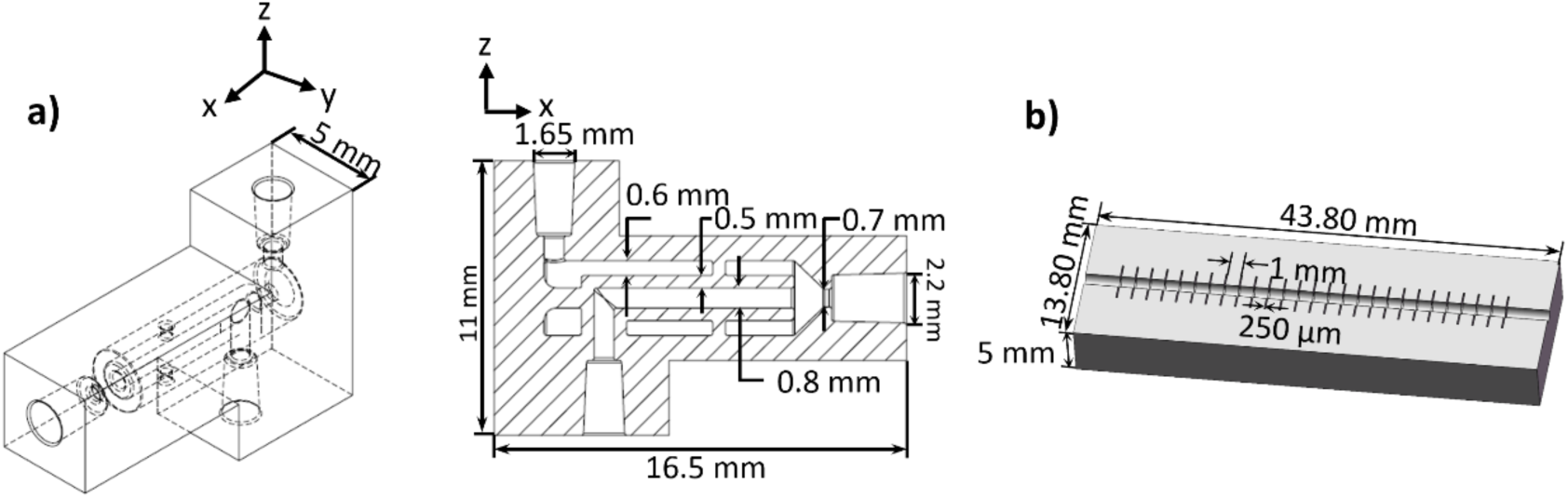
(a, b) The geometry of 3D printed microfluidic device(a) and photomask (b).

**Figure S8.**
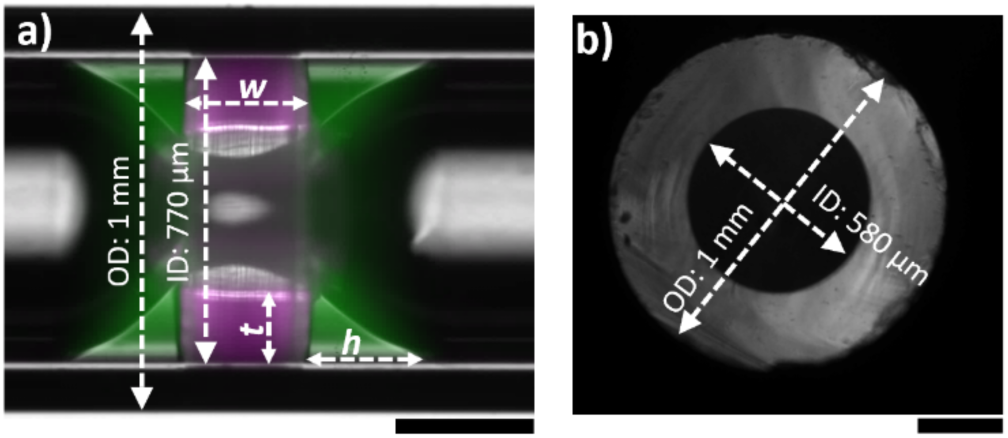
Experimental estimation of torodrop volume. (a) Geometric parameters measured from merged optical microscopy images. (b) Microscopy image showing the cross-section of the glass capillary. Scale bars represent 250 μm.

**Note S1. Numerical simulation of torodrop formation**

A two-phase, two-dimensional(2D) axisymmetric model was established to systematically investigate the torodrops formation. The effects of hydrogel ring inner radius (*r*), distance between adjacent hydrogel rings (*D*), and the thicknesses of the aqueous phase surrounding the hydrogel ring (*t*x and *t*y) were evaluated within a fixed simulation time **(Figure S1a**). Experimentally, the bulk aqueous solution was largely removed from the capillary during the unloading step, leaving a thin layer surrounding the hydrogel ring. Accordingly, thin aqueous layers were assumed in simulations (*t*_x_, *t*_y_ < 0.5 * *w*, where *w* is the width of the hydrogel ring) (**Table S2**). Because the hydrophilic glass capillary surfaces preferentially retained aqueous solution, *t*_x_ was assumed to be greater than *t*_y_. To mimic experiment conditions, simulations were performed with *t*_x_ < 100 μm and *t*_y_ < 60 μm while *r* and *D* ranged from 100 μm to 250 μm and 200 μm to 800 μm, respectively.

At computing time *T_sim_*=0, each hydrogel ring was surrounded by aqueous solution. It gradually spread over the hydrophilic capillary and hydrogel rings’ surfaces, achieving a minimum interfacial energy state within a short time.^1^ During the spreading process, the aqueous solution films on neighboring hydrogel rings and the capillary merged, resulting in the formation of a large single pinned drop when *D* = 200 μm and 400 μm. However, increasing *D* prevented the aqueous solution from merging, leading to two independent torodrops at *D* = 600 μm and 800 μm. This behavior was observed for all hydrogel ring inner radii investigated (*r* = 100 μm, 200 μm, 150 μm, and 250 μm) (Figure S1b). In all cases, the torodrops remained within the nanoliter volume range. The volume of merged single-pinned drops was approximately the sum of the two independent torodrops volumes (Figure S1c).

**Note S2. Experimental estimation of torodrop volume**

The thickness (*t*) and width of the hydrogel rings (*w*), distance between adjacent hydrogel rings (*D*), and inner diameter of the glass capillary were measured from merged optical microscopy images (**Figure S8a**) using ImageJ software. The measured inner diameter of the capillary (770 μm) was larger than its nominal inner diameter (580 μm), which was attributed to the cylindrical lens effect of the curved glass wall (Figure S8b). Refraction through the circular capillary wall magnified the apparent inner boundary, resulting in an overestimation of the capillary diameter in the microscopy images. In contrast, the measured *D* (around 1.0 mm) agreed well with the photomask design, indicating minimal distortion in the lateral dimensions. Therefore, a correction factor of 0.75, calculated as the ratio of the nominal inner diameter to the measured inner diameter (580/770), was applied to all measured hydrogel thickness values.

To estimate torodrop volume (*V*_t_), we mathematically model the geometry as a solid revolution comprising a central annular cylinder and two lateral revolved spandrels. The cross-section of the central region is defined as a rectangle of width (*w*) and height (*t*), located at a radial distance (*r*) from the central axis of rotation. The volume of central cylindrical (*V*_center_) segment is calculated as *V*_center_ = *πwt*(2*r* + *t*). Because the hydrogel is only partially occupied by water, the aqueous volume retained within the hydrogel ring (*V*_in_) was estimated by multiplying the geometric volume of the central cylindrical segment by the hydrogel water volume fraction (φ): *V*_in_ = *V*_center_∅. The *ϕ* was assumed to 0.40, corresponding to the water fraction in the precursor solution.

The lateral edges of torodrop are modeled by defining a concave spandrel, which is generated by subtracting a quarter-elliptical region from the bounding right triangle of width (*h*) and height (*t*). This ellipse is constrained to be tangential to both the top horizontal edge and the inner vertical boundary. The cross-sectional area of the resulting spandrel is given by 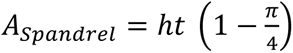. To determine the volume generated by revolving these lateral spandrels, we applied Pappus’s Centroid Theorem. By subtracting the first moment of area of the quarter-ellipse from that of the bounding rectangle, the local vertical centroid (y_c_) of the spandrel relative to its base is determined to be 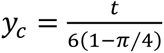. The volume of a single revolved lateral spandrel (*V*_s_) is thus the product of its area and the distance traveled by its centroid, *V*_s_ = 2*π*(*r* + *y_c_*)*A_spandrel_*, which simplifies to 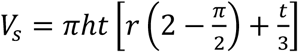. Thus, the *V*_t_ is calculated as 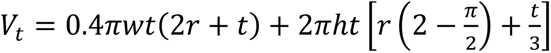.

The geometrical parameters and calculated torodrop volumes are summarized in **Table S3**.

**Note S3. Peclet number (*Pe*) calculation**

*Pe* number is defined as the ratio of diffusive time to convective time. 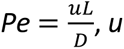 is the fluid velocity, L is the characteristic length, D is the diffusion coefficient.

The length and diameter of LabCap are 45 mm and 0.58 mm. The target solution was loaded into the capillary about 2 s by capillary force, respectively. The calculated average loading velocity *u* is 22.5 mm/s. As calculated, the *Pe* number was 6.5×10^4^ during capillary-driven sample loading, assuming a diffusion coefficient *D* = 10^−10^ m^2^/s, characteristic length *L* is 0.29 mm. When the target solution was injected into capillary using syringe pump at a flow rate of 10 μl/min, the corresponding *Pe* was 1830. In both cases, the high *Pe* value (≫ 1) suggested the target transport along the capillary is strongly convection-dominated. Consequently, sample loading under these conditions suppresses the formation of a diffusion-limited depletion zone, ensuring the uniform delivery of target molecules, whether loading is driven by capillary action or syringe-pump injection.

**Table S1.** Cost estimation for the establishment of LabCap.

| Item | Material cost | Cost (LabCap) |
| --- | --- | --- |
| Glass capillary | 80 €/ 500 capillary | 0.16 € (one capillary) |
| Microfluidic device | 2 mL Anycubic Aqua clear resin | 0.1 € (per device) |
| Photomask | 1 mL Phrozen Aqua Gray resin | 0.1 € (per device) |
| Silicon tubing | 10 meters/ 15 € | 0.03 € (2 mm) |
| PEGDA (Mw 575) | 80 €/ 100 mL | 0.01 € (per cycle) |
| PEG (Mw 200) | 60 €/ 1 L | 0.01 € (per cycle) |
| PI | 100 €/ 500 mL | 0.01 € (per cycle) |
| Total cost (per device) |  | 0.42 € |

**Table S2.**
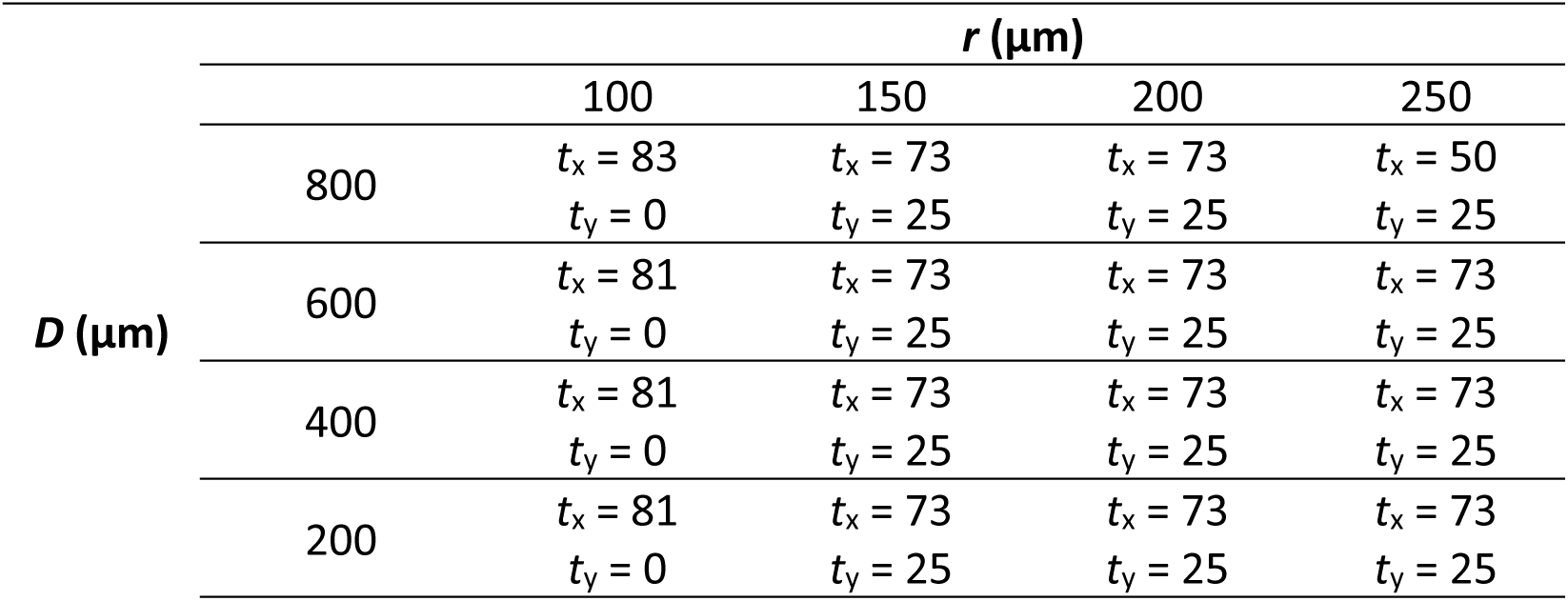
Numerical simulation parameters for the aqueous phase thickness (*t*_x_ and *t*_y_, µm) surrounding the hydrogel ring.

**Table S3.** Experimental measured geometrical parameters of hydrogel rings and calculated torodrop volumes.

| $Q_{\text{PEG}}:Q_{\text{PEGDA}}$ | 1 : 1 | 2 : 1 | 4 : 1 | 6 : 1 |
| --- | --- | --- | --- | --- |
| $t$ ($\mu\text{m}$) | 132.2 | 94.6 | 72.9 | 50.1 |
| $r$ ($\mu\text{m}$) | 157.8 | 195.4 | 217.1 | 239.9 |
| $h$ ($\mu\text{m}$) | 270.4 | 218.9 | 221.6 | 152.6 |
| $R$ ($\mu\text{m}$) | 290 | 290 | 290 | 290 |
| $V_t$ (nL) | 47.4 | 32.3 | 25.9 | 15.8 |

**Table S4.**
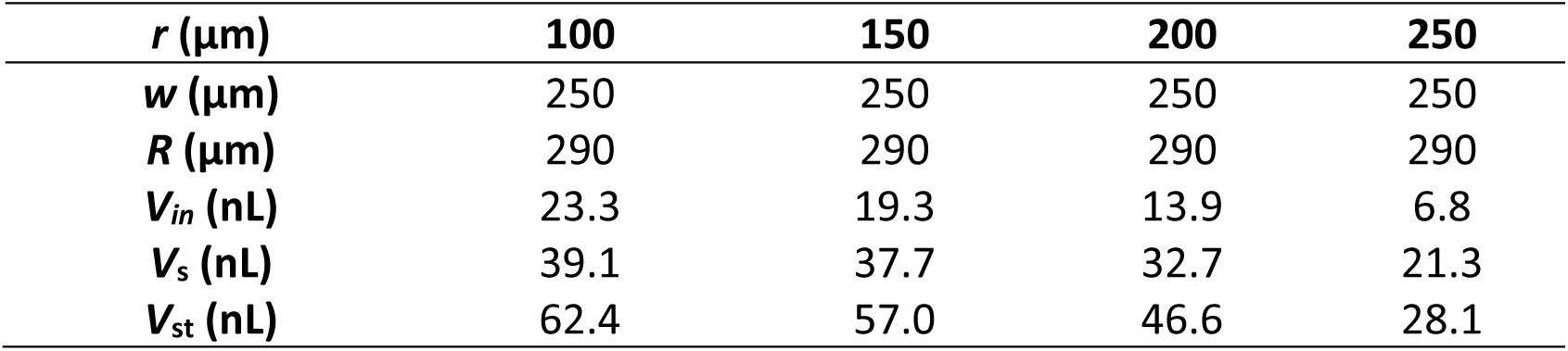
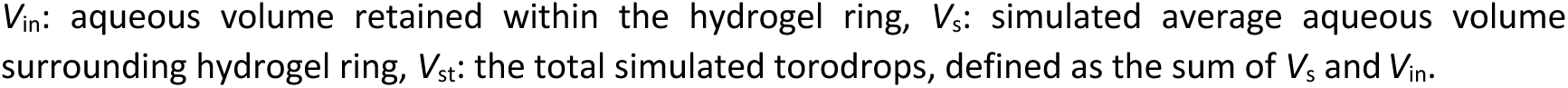
Simulated volumes of torodrops generated using hydrogel rings with varying inner radii (*r*) at a fixed distance (D) between adjacent hydrogel rings of 800 μm.

**Table S5.** Reagent volume estimation for the ELISA performance on LabCap platform using different incubation protocols.

| Reagent | Medium exchange ($\mu\text{L}$) | Static incubation ($\mu\text{L}$) |
| --- | --- | --- |
| Streptavidin | 96 | 12 |
| Capture antibody | 96 | 12 |
| Blocking buffer | 96 | 12 |
| Detection antibody | 96 | 12 |
| Sample | 96 | 12 |
| QuantaRed | 12 | 12 |
| Total wash buffer | 1000 | 1000 |

